# HyLight: Strain aware assembly of low coverage metagenomes

**DOI:** 10.1101/2023.12.22.572963

**Authors:** Xiongbin Kang, Wenhai Zhang, Xiao Luo, Alexander Schönhuth

## Abstract

Different strains of identical species can vary substantially in terms of their spectrum of biomedically relevant phenotypes. Reconstructing the genomes of a microbial community at strain level poses major methodical challenges, because relative frequencies of individual strains match the rate of sequencing errors, which hampers the identification of their characteristic genetic variants. While next-generation sequencing (NGS) reads are too short to span complex repetitive regions, the considerably longer third-generation sequencing (TGS) reads are affected by larger sequencing error rates or are just significantly more expensive. Suppressing TGS coverage to limit costs implies losses in terms of the accuracy of the assemblies. Therefore, existing approaches have remained fragmentary: all approaches presented so far agree on losses in strain awareness, accuracy, possibly excessive costs of the assemblies, or combinations thereof.

We present HyLight as, to the best of our knowledge, the first metagenome assembly approach that is not affected by any of the aforementioned drawbacks. In experiments, we demonstrate that HyLight assemblies are strain-aware, contiguous, contain little errors, and because operating on low coverage TGS data of the cheap kind, come at drastically reduced costs. HyLight implements hybrid assembly, which exploits the complementarity of TGS and NGS data. For unifying the two types of data, HyLight utilizes strain resolved overlap graphs (OG), which support the accurate reconstruction of the individual members of microbial communities at strain level: HyLight outperforms existing approaches in terms of strain identity preserving sequence by on average 25.53% (across all experiments / data sets: first quartile: 21.53%, median: 26.81%, third quartile: 31.98%), achieving near-complete strain awareness on many data sets. In summary, HyLight appears to implement the first protocol that delivers assemblies that are strain-aware, contiguous and accurate in combination.

## Introduction

Metagenomics has revolutionized our understanding of microbial diversity and functional potential in various environments by sequencing the collective genomic material of microbial communities in many niches. The advent of high-throughput sequencing technologies (NGS) has made metagenomic research increasingly accessible, providing valuable insight into complex microbial ecosystems (1; 2; 3).

However, the assembly of metagenomic data poses significant challenges due to the presence of multiple strains, high levels of genetic diversity, and varying abundances of different organisms within a community (4; 5). Strains that appear at only low abundance in a metagenome, whose discovery however is crucial in the context of understanding the environment under investigation, stress the practical relevance of these challenges.

In further detail, strains can display significant variations in their interactions and environmental impact (6; 7). Strains can also differ in clinically relevant aspects such as medication resistance, virulence, and host-microbiome interactions, which can have serious consequences on human health (8; 9; 10; 11). For these reasons, it is crucial to identify genomes at strain resolution. This explains why *strain aware metagenome assembly* has become the currently driving technical and methodical challenge (12; 13).

Next-generation sequencing (NGS), where Illumina sequencing technology is the leading option, is extensively utilized in metagenomics research for being culture-independent, and its combination of high throughput, accuracy, and cost-effectiveness (14). However, NGS produces short reads typically ranging from 35 bp to 700 bp in length (15). The short length of NGS reads induces obvious difficulties in spanning complex genomic regions (16; 17), which leads to fragmented genome assemblies (18; 19; 20). This limitation becomes particularly concerning when processing metagenomic samples of greater complexity, characterized by harboring diverse strains (19). In summary, the reconstruction of both strain-level resolved and complete genomes of individual organisms from metagenomes is too challenging when using NGS alone.

Third-generation sequencing (TGS), such as Pacific Biosciences (PacBio) and Oxford Nanopore Technologies (ONT), holds the promise to overcome these challenges. Because TGS reads are significantly longer, they can span complex regions, which makes it possible to distinguish between related organisms (15; 21; 22; 13). On the other hand, however, TGS is subject to considerably higher error rates and substantially larger costs. In contrast to NGS platforms with sequencing error rates below 1%, even PacBio CLR and ONT reads that still exhibit greater error rates (ranging from 2 to 15%) imply substantially larger costs because of the elevated coverage required to compensate the errors (21).

The latest generations of PacBio HiFi or ONT Q30+ type reads do not alleviate this issue, because their lower error rates (approximately 1%) are put into perspective by both their substantially shorter length (still exceeding 10 kb, but being up to 4-8 times shorter than PacBio CLR or ONT reads (21)), their lower throughput in general, and their substantially larger costs (21). An additional detail is the fact that the process of converting subreads into HiFi reads using the CCS algorithm beyond the monetary efforts also requires substantial computational investments; for example, it takes 10 000 CPU hours to process the output of a single SMRT cell concerned with a data set of 7 million reads (21; 23). In summary, PacBio HiFi or recent generations of ONT type reads mean practical convenience that only generously equipped laboratories can afford. The vast majority of sequencing laboratories, however, keeps depending on NGS on the one hand, and cheaper hence more erroneous (but in case of PacBio CLR and ONT still the longest!) types of TGS reads.

In summary, sole application of TGS in metagenomics requires to substantially raise expenses: either to increase coverage (as for PacBio CLR and ONT) or because of employing more sophisticated sequencing protocols (PacBio HiFi or ONT reads of latest generations). In addition, these more sophisticated protocols entail substantial additional requirements in terms of computational resources (24; 21; 25). *The great majority of labs worldwide cannot employ TGS (of whatever kind) alone for strain-level metagenome assembly, either because coverage remains too low or expenses become excessive*.

Because of this mix of theoretical and practical challenges, only very few methods have been dedicated to strain aware metagenome assembly so far. Current state-of-the-art approaches are StrainXpress (12) on the one hand, which focuses on NGS, and Strainberry (13), on the other hand, which focuses on TGS, as the only two approaches that have decidedly targeted at *strain aware* metagenome assembly. As we will demonstrate, however, neither StrainXpress nor Strainberry are able to generate assemblies of the kind of quality that we are envisioning. While StrainXpress assemblies remain too fragmented because of the short length of NGS reads, Strainberry requires elevated coverage rates for the TGS reads, which, as repeatedly pointed out, is expensive. Note that our particular vision here is to present a strain aware metagenome assembly approach whose demands in terms of TGS read coverage are low and which does not require expensive technology. We conclude that sole application of either only NGS, on the one hand, or only TGS, on the other hand, for the purposes of strain aware metagenome assembly is confined to a handful of more amply budgeted laboratories (25). This hampers the routine and widespread implementation of strain-aware metagenome analyses in disease research.

It is important to realize that both StrainXpress and Strainberry employ techniques that had hitherto never been considered for metagenome assembly. On the one hand, StrainXpress employs overlap graphs instead of de Bruijn graphs (DBGs) where the latter ones have been the (by far) predominant data structure paradigm employed for NGS based assembly. The crucial insight here is the fact that usage of OG’s effectively aids in spanning complex repetitive regions, because OG’s do not require to chop reads into *k*-mers. As a consequence, genetic linkage of strain specific variants becomes evident. This makes it possible to extend contigs across regions that remain “strain-specific variant deserts” when operating with NGS reads or de Bruijn graphs.

Strainberry, on the other hand, initially employs Metaflye (26) to assemble the TGS reads into contigs, and subsequently aligns the long reads with the contigs. Based on these alignments, Strainberry calls SNP’s where contigs serve to provide auxiliary reference coordinates. The resulting scenario provides the basis for solving the minimum error correction problem to phase reads into haplotypes. The source of inspiration is earlier work that proved that modeling haplotype separation as instances of the minimum error correction problem was useful (27; 28; 29). As a novelty, Strainberry applies this procedure in the frame of an iterative protocol: the iteration ends when separation does no longer reveals novel haplotypes; eventually strain-resolved contigs are assembled using Wtdbg2 (30).

In summary, both StrainXpress and Strainberry, as the sole approaches to address *strain aware* metagenome assembly in a decided manner, employ novel, so far non-conventional techniques. It appears that methodical novelties, which are rather surprising from a historical perspective, are crucial for successful strain aware metagenome assembly: as repeatedly pointed out, application of standard techniques runs into either larger costs, lower assembly quality, or both.

In this paper, we would like to present an alternative strategy that is based on an unusual protocol, on the one hand, and on unusual techniques as elements of the protocol, on the other hand. We envision that our approach is the first one that enables the majority of laboratories to perform strain aware metagenome assembly without incurring costs that are excessive. Beyond our strategy being a considerably cheaper alternative, our strategy also outperforms all prior state-of-the-art approaches by its assemblies being largely superior over the more expensive strategies. So, also laboratories that can afford the practical convenience of expensive protocols may be interested in integrating our approach into their practice: in terms of the quality of the metagenome assemblies our strategy arguably offers choices that are superior also from a comprehensive perspective.

The basis for our strategy is hybrid assembly. Hybrid assembly synthesizes the advantages of short and long reads. Importantly, short and long reads complement each other perfectly: while NGS reads are accurate and short, TGS reads are long but inaccurate. Synthesizing them, when done well, yields assemblies that rely on long and accurate pieces of sequence as a basis, which constitutes the optimal scenario. The practical feasibility of hybrid assembly is established by the fact that the vast majority of sequencing laboratories are equipped with (Illumina type) NGS platforms and TGS platforms of the cheaper types.

So. pursuing hybrid assembly strategies perfectly caters to our vision of offering strain-aware metagenome assemblies “for everyone”. In addition, importantly, our protocols are not very affordable, but also lead to assemblies whose quality substantially pushes the limits of metagenome assembly from a global perspective.

The general benefits of hybrid assembly, namely the accuracy of the assemblies and the inexpensiveness of the supporting data have already been noticed in prior work. Despite these general benefits, however, all current state-of-the-art hybrid assembly approaches that can be applied for metagenomes (31; 32; 33; 34) operate at the species level, as the finest taxonomic resolution. Note that all of them have been widely applied in studies that require metagenomic assembly as an essential step (35; 36; 37; 38; 39).

To understand why the state-of-the-art can only deliver species-resolved assemblies, one needs to have a closer look at the methodologies that underlie the current hybrid assembly approaches, both when specializing in metagenome assembly or when used for other experimental scenarios.

Hybrid assembly broadly falls into two categories: *short-read-first* and *long-read-first* approaches. *Short-read-first approaches* assemble short reads first, and then scaffold the short read based contigs using the long reads. An important observation is that short-read assembly is dominated by de Bruijn graph (DBG) based approaches, due to the fact that DBG’s have been found to have substantial advantages when processing NGS reads. Vice versa, *long-read-first* approaches use long reads as the foundation for their assemblies, and treat short reads as an auxiliary source of data. This implies in particular that long-read-first approaches never assemble short reads in their own right. Rather, they make use of the short reads to correct the long read based assemblies.

There are two principled subcategories of long-read-first approaches. Either, they assemble the raw long reads, and subsequently align the short reads against the long read contigs to eliminate the (usually numerous) sequencing induced errors from the long read contigs (40). The second subcategory, on the other hand, is characterized by using the short reads for eliminating the errors from the raw long reads, and to assemble the long reads after their short read assisted correction (41).

The first striking observation is that all current state-of-the-art in hybrid metagenome assembly (33; 31; 34; 32) are short-read-first approaches. In particular, all of them make use of DBG’s to assemble the short reads in the first step of their protocols.

In a bit more detail, Opera-MS (33) first uses an established short-read metagenome assembler like MegaHit (42), MetaSPAdes (43) or IBDA-UD (all of which are DBG based (44)) to assemble the short reads. Subsequently, both short and long reads are mapped against the short-read contigs to obtain coverage and linkage information for the contigs, and to construct an assembly graph based on that information. Contigs are further hierarchically clustered using a distance measure that reflects the distance between the contigs in terms of their distance in the assembly graph. Apart from certain details, each cluster is supposed to collect the contigs of one species. Finally, contigs within clusters are scaffolded using Opera-LG, which was designed to work for the hybrid assembly of isolated genomes.

HybridSpades (31) constructs an assembly graph from the short reads using SPAdes (45), by removing bulges, tips and chimeric edges. By mapping the long reads against the resulting assembly graph, it generates read paths, and further closes gaps in the assembly graph by using the consensus of the long reads that span the gaps. Then, by extending a technique that addresses to grow read paths by trying “extension edges” to incorporate long read paths, it resolves repeats.

MetaPlatanus (34) computes contigs by constructing a DBG, and subsequently corrects contigs based on coverage considerations relating to both short and long reads, and also untangles “cross-structures” from such considerations. DBG’s are re-constructed iteratively, where in each iteration the improved contigs serve as the basis for constructing the DBG, and subsequent correction of contigs yields new contigs. MetaPlatanus then scaffolds the resulting contigs using long read links, and, possibly, re-constructs the DBG another last time. Eventually, scaffolds are binned, where each bin is supposed to refer to a particular species. Within bins, gaps are closed, edges are extended, and possibly unused short reads are employed for aiding in that. In an ultimate step, gaps are closed and scaffolds are polished using techniques that address the generation of assemblies for isolate genomes.

Unicycler (32), like HybridSPAdes, uses SPAdes (45) for assembling the short NGS reads. Like MetaPlatanus, it employs coverage considerations to refine the resulting assembly graph. Subsequent integration of long reads then points out paths in the refined assembly graph. “Bridges” that reflect resulting new links in the assembly graph are ranked by quality (measured in terms of coverage, for example), and then are applied in decreasing order relative to their quality, which establishes the paths that are supposed to be real. The resulting assemblies are finally polished by re-aligning the short reads against the selected paths using Bowtie (46), as a standard short-read mapper.

As a summary of the state-of-the-art that either reflects the current standards in metagenome assembly in general, and the current standards in hybrid metagenome assembly in particular, we count StrainXpress and Strainberry, on the one hand, and Opera-MS, HybridSPAdes, MetaPlatanus and Unicycler, on the other hand. In the following, we will compare our approach with these six prior approaches, because these six approaches appear to have established the state-of-the-art with respect to all aspects of interest here.

In comparison with the earlier hybrid assembly approaches, we suggest a strategy that neither reflects a “short-read-first” nor “long-read-first” approach. Instead, we suggest to compute assemblies from the long reads and the short reads, where short reads assist in the assembly of the long reads and vice versa; that is, we treat both long and short reads as both primary assembly and auxiliary data (note that we avoid to compute assemblies that can be foreseen to be redundant, to avoid unnecessary runtime leaks). Finally, we merge the two assemblies into a unifying set of (scaffolded) contigs. The idea to not distinguish the two types of data into fundamental assembly data on the one hand, and auxiliary data on the other hand, appears to be novel in comparison with all (both generic and specific metagenome) hybrid assembly approaches. From this perspective, our approach can be viewed to establish a third and novel category, which one could refer to as *”cross hybrid”* (or, alternatively, “mutual support”) approach.

From another perspective, our approach is also novel insofar as it is the only approach that makes use of overlap graphs, instead of DBG’s, as a unifying data frame for all sources of data involved. In that respect, our approach draws inspiration from StrainXpress, which demonstrated that OG’s could also be used in an advantageous way when it comes to distinguishing between similar strains of the same species. To understand this better, we recall that the construction of DBG’s implies to chop reads into k-mers. The artificial shortening of the reads leads to significant losses in terms of information with respect to genetic linkage of strain specific variants (in particular, one can no longer trace linkage of variants at distance larger than *k* when employing *k*-mers (47)). Unlike DBG’s, OG’s preserve the haplotype (strain) identity of the reads to a maximum degree, which we systematically exploit also here. On a side remark, note that the employment of OG’s also lead to superior strain awareness in viral quasispecies assembly (48; 49).

### Summary of Contributions

In summary, the novelties we suggest are as follows.

1. We suggest the first hybrid metagenome assembly approach that is strain aware.
2. In all earlier approaches, either short reads or long reads were used as fundamental assembly data where the other type of reads was used as auxiliary data. We suggest the first hybrid assembly approach that makes use of both long and short reads as both fundamental assembly and auxiliary data, and so is *”cross-hybrid”* in nature.
3. We suggest the first hybrid assembly approach in which overlap graphs are used to capture effects relevant for the assembly of the short reads. A particular feature is the employment of “contig OGs”, as a rather unusual concept in genome assembly.
4. We suggest the first metagenome asssembly approach in general that is strain aware without requirements in terms of long read coverage or sophisticated sequencing protocols that tend to be excessive in terms of costs.
5. Last but not least, we suggest a metagenome assembly approach that is superior over all prior approaches in terms of assembly quality. Arguably, by its results, HyLight considerably pushes the limits of the possible in strain aware metagenome assembly, all in terms of strain awareness, contig length (contiguity), and in terms of the accuracy of the contigs.

In the following, in Results, we will present the workflow of our approach as well as all details required for understanding it from a larger perspective. Subsequently, we will present the experiments that confirm the novelties and improvements as pointed out above. As usual, we will discuss our results in comparison with the hypotheses just raised. In Online Methods, we will provide all the details that establish the full reproducibility of our approach.

## Results

In the following, we first briefly discuss the workflow. Details that are necessary to fully reproduce the workflow from a conceptual point of view are provided in Methods. Subsequently, we present experiments on both simulated and real data, which provide evidence for the benefits of our approach, as listed towards the end of the Introduction.

HyLight’s major innovation lies in the construction of a strain-resolved overlap graph (OG) as input for assembling long reads, correcting contigs, and clustering and assembling short reads, ultimately achieving strain-aware assembly. The general steps are as follows: firstly, short reads are used to correct long reads, resulting in high-quality long reads. Then, we use the corrected long reads to construct a strain resolution OG, so that the assembled long-read contigs have high accuracy and strain resolution. However, due to the low sequencing coverage or high error rate in some regions of the long reads that are not well corrected, some strains or regions cannot be well reconstructed. At this point, we need to use short reads to assist with assembly. First, we align short reads to the long-read contigs, then untangle erroneous overlaps to obtain a strain-aware OG of short reads. However, this OG is not used for assembly, but to filter out short reads belonging to strains that have already been assembled. After filtering out short reads of the assembled strains, we use the remaining short reads to construct a short reads OG and continue assembling them. Finally, we collect the contigs generated by long reads and short reads together, construct a contig OG, and further extend contigs.

### Workflow

Please see Figure 1 for a schematic of the workflow. The workflow proceeds in two axes: one for assembling the long reads (see left branch in Fig. 1) and one for assembling the short reads (right branch in Fig. 1). Assemblies of long and short reads are merged in a final step (see bottom of Fig. 1).

**Figure 1.**
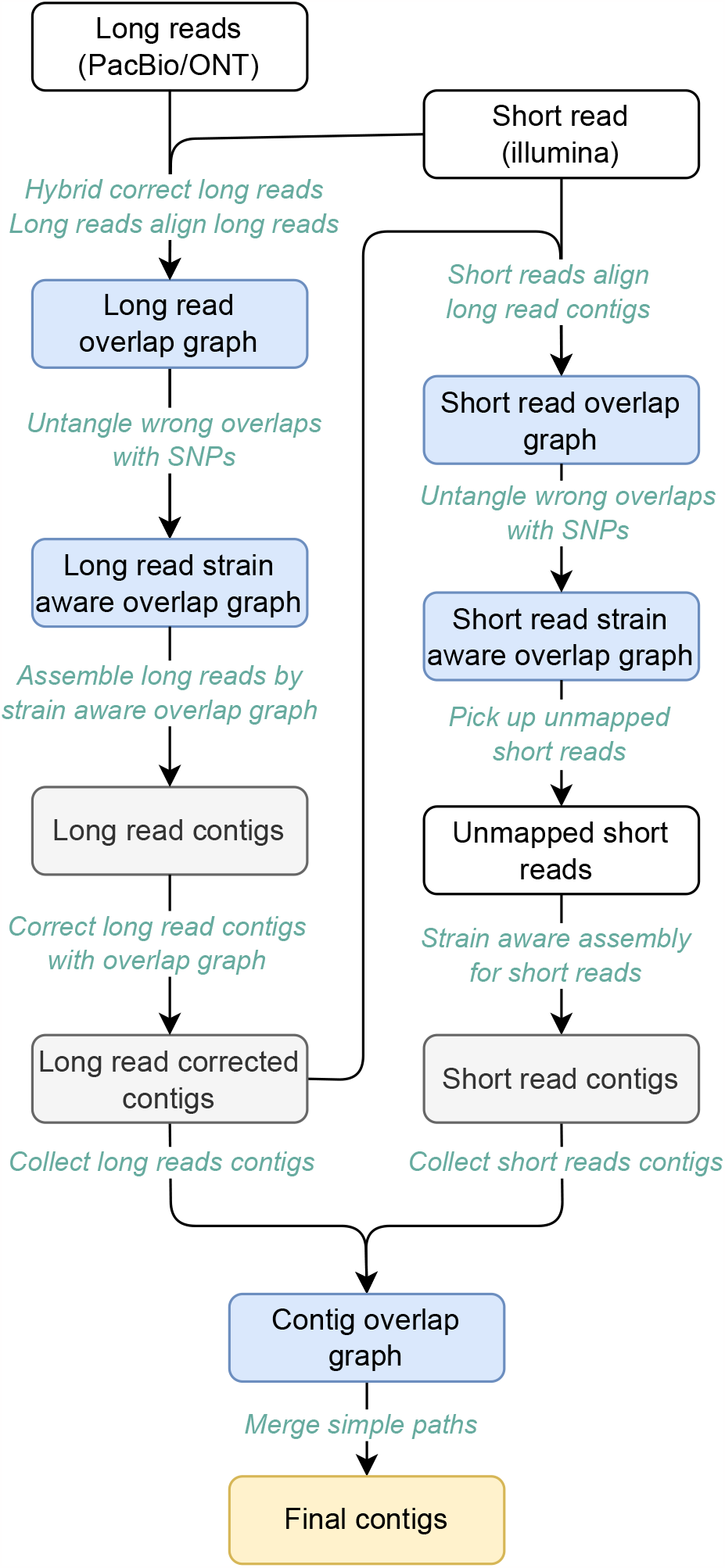
Workflow of HyLight. The input data consists of two fastq files, long reads and short reads. The output is a fasta file containing the assembled contigs. The overall procedure can be divided into three primary steps. Firstly, strain-resolved OG is conducted to assemble long reads. Subsequently, another OG is established to assemble short reads. Lastly, a contig OG is established to extend the contigs obtained from the assembly results of long and short reads, culminating in the generation of the final master contigs.

In a brief summary, HyLight performs the following items. First, it corrects long reads using short reads, which turns the raw TGS reads into polished, error-free long reads. The resulting polished long reads then are the basis for constructing a strain-resolved in error-removed, polished

To provide an overview of the workflow, this section offers a high-level description. For detailed descriptions of all methodical steps involved, please refer to the Methods section. Figure 1 illustrates the overall workflow of HyLight. As outlined previously, HyLight comprises three main modules.

#### First module: long read assembly

The main purpose of this axis is to compute strain-aware, error-free contigs from the long reads.

1. Long reads are corrected using short reads using FM-index and de Bruijn graph based techniques, as implemented in FMLRC2 (50), which has been shown to outperform other methods in recent benchmark studies (51; 52).
2. An overlap graph is constructed from the corrected long reads. We make use of the (widely popular) Minimap2 (53) to compute the necessary overlaps.
3. We identify overlaps that connect long reads from different strains by inspecting SNP patterns, and we remove edges in the overlap graph that reflect the connection of long reads from different strains. See Figure 2 for an illustration. The result is an overlap graph of the long reads that consists of connected components each of which contains long reads from only one particular strain. So, each connected component in the overlap graph now reflects a collection of reads drawn from one haploid genome.
4. We assemble the long reads based on the resulting strain-aware overlap graph using Miniasm (54), which is a long read assembler that addresses to assemble haploid genomes from long reads. The result are contigs each of which stems from one particular strain.
5. We re-align the long reads against the resulting strain-aware contigs.
6. Based on the re-alignment, we establish a second, improved version of an overlap graph for the long reads, which now reflects a *strain-aware overlap graph* of the long reads.
7. Using this improved, strain-aware overlap graph, we remove errors that have remained in the long reads using Racon (55).

**Figure 2.**
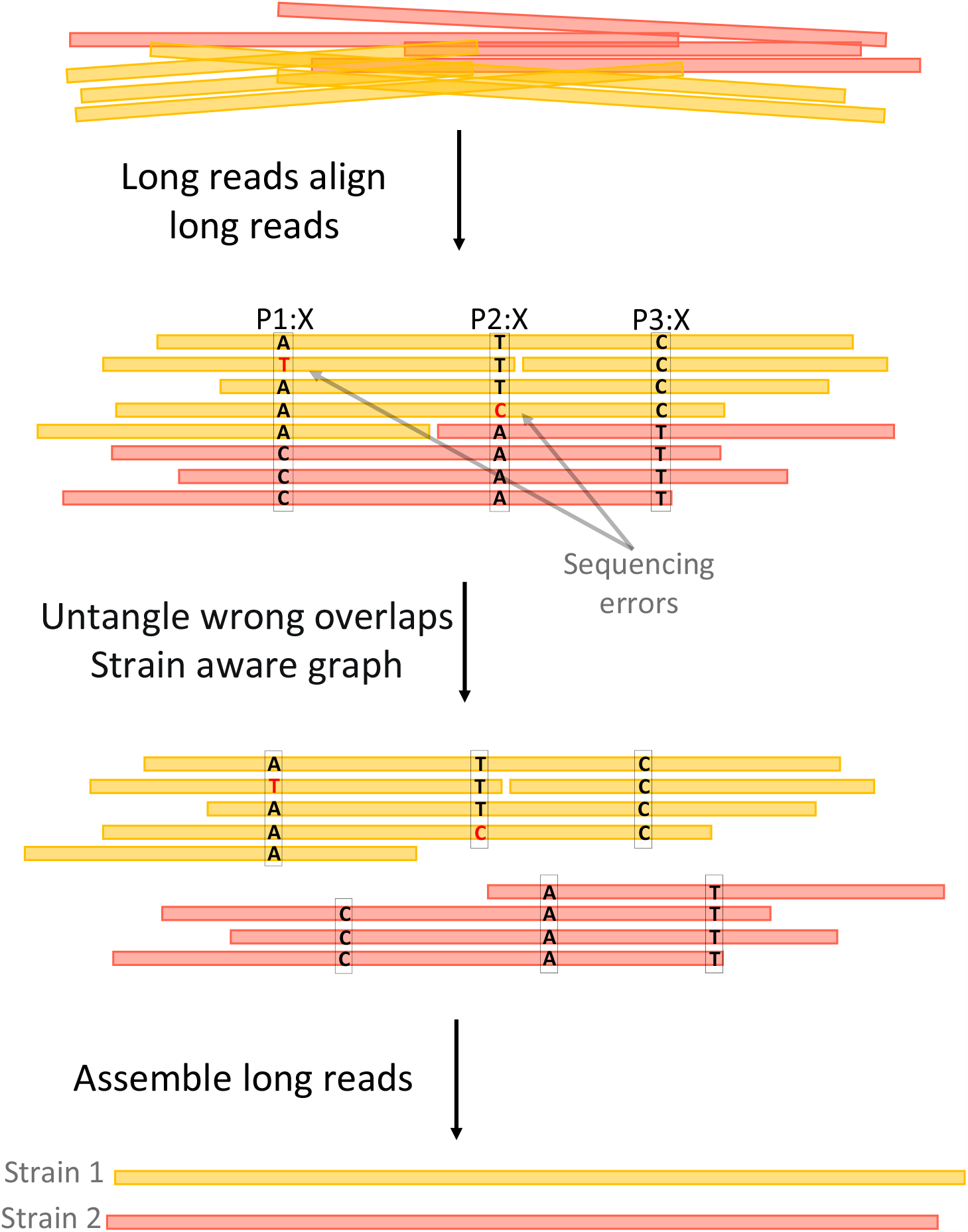
Assemble long reads. The long reads used here are hybrid corrected reads. The distinct colors of the reads indicate their respective strain origins. The objective of this workflow is to leverage SNP information to filter out incorrect overlaps, selectively retaining overlaps between reads originating from the same strain. This enables strain aware assembly to be performed effectively.

As for 7.), note that Racon has not been designed to operate in a strain-aware manner. If one fed the original, strain-unaware overlap graph to Racon, it would “overcorrect” contigs by mistaking true, strain-specific variation for errors and eliminating them. This would mask strain-specific variation hence prevent the reconstruction of genuinely strain-specific sequence. One can consider the application of Racon to only strain-aware overlap graphs an insight that is crucial for computing both error-free *and* strain-aware long read based assemblies.

#### Second module: short read assembly

We recall that it is a general objective to establish a workflow that caters to *low long read coverage* to avoid unnecessary and possibly unaffordable costs. Therefore, the main purpose of this axis is to assemble the (likely high coverage, because cheap) short reads in their own right, and use the assemblies to fill gaps in the long read based assembly, or even identify additional strains from the resulting contigs.

1. We align the short reads against the strain-aware, error-free contigs, as the output of the long read axis (first module), using Miniasm (54).
2. The alignment of short reads with long read contigs gives rise to an overlap graph of the short reads.
3. Analogously to the long read axis, we inspect SNP patterns in the overlap of the short reads. Based on the SNP patterns, we identify overlaps of short reads that reflect to connect two short reads from different strains. See Figure 3 for an illustration.
4. As a result, one is now able to identify short reads whose SNP patterns contradict their initial alignment with the long read contigs. The insight is that breaking up overlaps between short reads all of which align with the same long read contig leads to several classes of short reads. Only one of the classes of reads truly agrees with their respective long read contig (yellow short reads in Fig. 3).
5. One concludes that short reads no longer having overlaps with short reads whose SNP patterns truly match those of their long read contigs, do not stem from the same strain as the long read contig against which they initially aligned (blue short reads in Fig. 3).
6. Further, one collects all short reads that did not align with any of the long read contigs (grey short reads in Fig. 3).
7. One discards all short reads whose alignments indicated full agreement with a long read contig (yellow in Fig. 3).
8. Subsequently, using StrainXpress (12) (which as discussed in the Intro specializes in the strain-aware assembly of short reads using OGs), one assembles all short reads whose alignments were not in full agreement with their long read contigs (blue in Fig. 3) or which had remained entirely unaligned with long read contigs (grey in Fig. 3). See the bottom of Fig. 3 for an illustration of the resulting assemblies.

**Figure 3.**
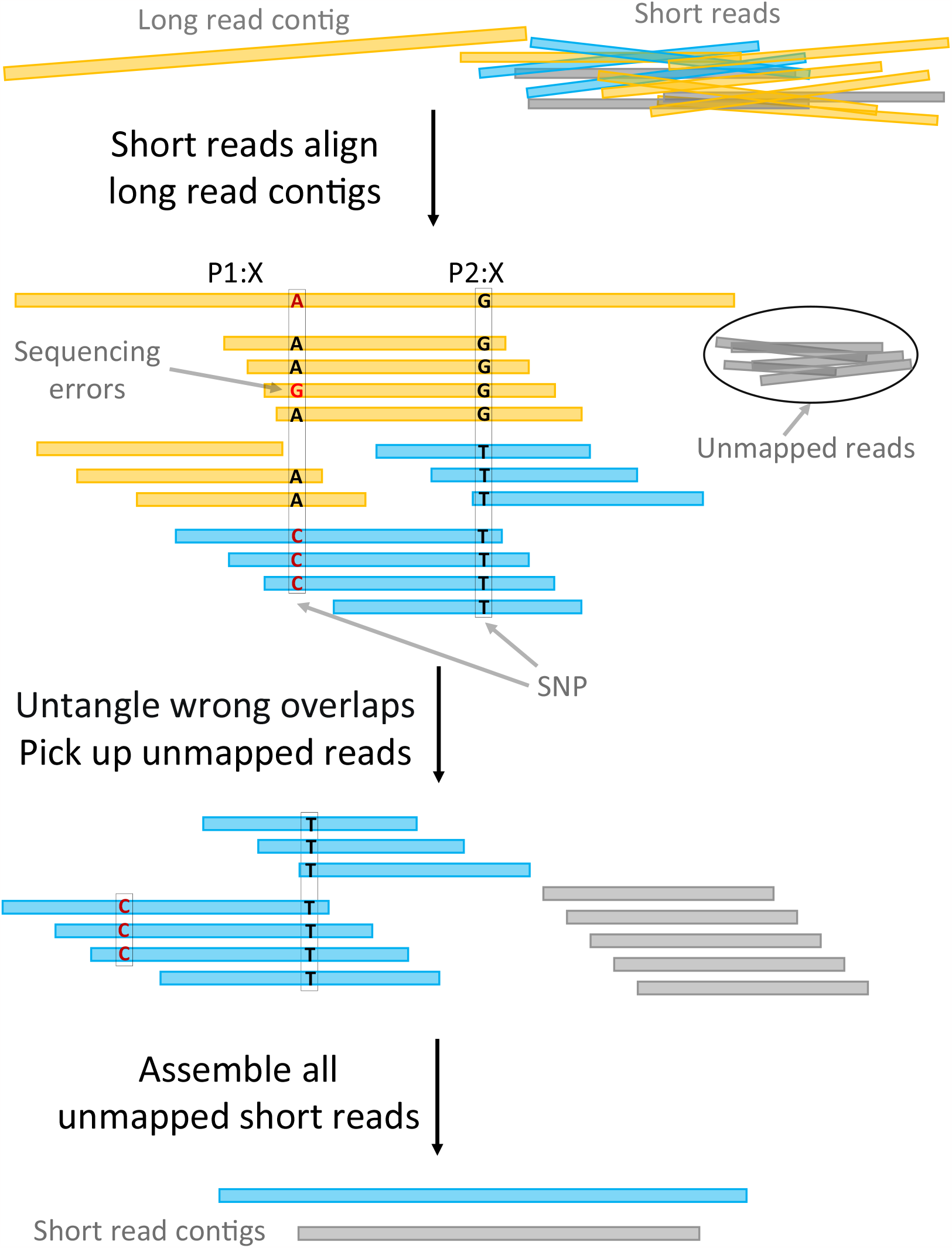
Assemble short reads. The distinct colors of the reads indicate their respective strain origins. The primary procedure consists of aligning the short reads to the contigs, establishing a strain-resolved OG, and then excluding short reads that align to regions already assembled into contigs. Subsequently, an OG is constructed to assemble the remaining short reads and reconstruct strains or regions that were not initially assembled.

As per the properties of StrainXpress, the result are strain-aware short read based contigs. As per the protocol we follow, all contigs refer to strain-aware metagenomic sequence not captured / spanned by any of the long read contigs from the first module.

#### Third module: merging long and short read assemblies

The purpose of this final module is to compute a unifying assembly that is as comprehensive as possible, as the final output of our approach.

1. One collects both long and short read contigs, as the output of the first and second module, and computes overlaps between them to establish an *encompassing strain-aware overlap graph*.
2. One identifies nodes in the OG through which only one particular path passes (”simple path”).
3. One extends contigs along the identified “simple paths”. The resulting extended contigs are the final outpu.

### Data & Experimental Setup

The experiments we discuss in the following refer to both simulated and real data.

The *synthetic data* we treat refer to scenarios that reflect different levels of complexity. In particular, we deal with data sets reflecting 3 *Salmonella* strains, and further 20 (low complexity), 100 (medium complexity) and 210 (high complexity) bacterial strains. Strains were retrieved from (56, DESMAN); see Supplementary Table S1 for full information on strain composition of data sets. All data sets were simulated using CAMISIM (57), which reflects a state-of-the-art and widely popular choice for generating metagenome sequencing data sets. Further, we also consider 6 “strain-mixing spike-in” data sets, which reflect spiking simulated reads from *Salmonella* strains into real data. This creates a real data scenario for which ground truth (in form of simulated reads) is available. See the Methods section for full technical details.

The *real data* are two microbial communities that reflect the current standard in terms of available real, *both TGS and NGS*, data with known ground truth. Both Bmock12 (a bacterial mock community) and NWC (a natural whey culture data set) have already been widely used in the evaluation of metagenome assembly approaches (52; 58; 13; 59; 12). For both data sets, reference genomes, Illumina, PacBio CLR and ONT reads are readily available. See again the Methods section for full technical details.

### Benchmarked Approaches

As discussed in the Introduction, the state-of-the-art when comparing hybrid metagenome assembly approaches that operate in a strain-aware manner, are first Strainberry, on the one hand, as the leading approach to deal with only TGS data and second StrainXpress, on the other hand, as the leading approach to only deal with NGS data. Hybrid metagenome assembly approaches that address strain awareness have not yet been presented before; here, we consider all state-of-the-art approaches to metagenome assembly that operate on the species level. These are HybridSPAdes (31), MetaPlatanus (34), Unicycler (32) and Opera-MS (33).

### Note on Metrics

In the following, we evaluate the performance in terms of metrics that are routinely computed by MetaQUAST V5.1.0rc1 (60). We particularly focus on “Genome Fraction” (GF), as a metric that refers to strain awareness (GF = 100.0 translates into full strain awareness), NGA50, as a metric deemed sufficiently reliable to measure contig contiguity, and (mismatch / indel) error rates as well as miassembled contig fraction (MC) to evaluate the quality of the contigs. See Methods for full details.

### Note on classification of approaches

We recall that the competing hybrid assembly primarily target at the accuracy and the length of the assemblies, and do not address strain awareness. Strainberry and StrainXpress, on the other hand, which use only TGS and only NGS data, respectively, primarily target at strain awareness, but suffer from more erroneous (Strainberry) or shorter (StrainXpress) assemblies due to the nature of the type of data they use as their input. HyLight is the only approach that targets at all of accuracy, contiguity and strain awareness.

Here, because of the different primary goals of the approaches, we would like to avoid to compare prior hybrid assembly approaches with prior non-hybrid approaches that focus on strain awareness. Therefore, in the following, we first compare HyLight with the prior hybrid assembly approaches, and, subsequently, in separate paragraphs, present a comparison of HyLight with Strainberry and StrainXpress.

#### Misassembled Contig Rate

To avoid repeating statements When comparing HyLight with Strainberry and StrainXpress in terms of misassembled contig fraction, we will not go into detail with respect to each of the data sets we run experiments on. As a *general trend* —which applies with no exception on any of the data sets—HyLight and StrainXpress considerably outperform Strainberry, which for (the solely NGS based) StrainXpress can certainly be attributed to the reduced length of the contigs, and for Strainberry can be attributed to being based on solely TGS data, which prevents the detection of misassemblies thanks to the accuracy of auxiliary NGS data.

### Experiments: Synthetic Data Sets

#### 3 Salmonella

This data set contains simulated reads from three distinct strains of *Salmonella*. The average coverage for Illumina (NGS) and PacBio (TGS) reads is 20X and 10X, respectively, reflecting a low-coverage TGS data scenario in particular, as intended. Despite the low number of strains (3), the high degree of similarity between them ensures that only approaches that are sufficiently strain aware are able to assemble them without confounding them. The data set serves as a test bed for evaluating basic properties of the benchmarked approaches. See Methods for full details.

##### Hybrid Assembly Approaches

See Table 1 for corresponding results. HyLight outperforms all other hybrid assembly approaches in terms of all relevant metrics. It covers 23.78% more strain sequence than the second best hybrid assembly approach (HyLight: 96.03 / MetaPlatanus: 72.25), missing out on only 4% strain-specific sequence. HyLight also dominates the other approaches in the other relevant categories, where improvements are to be measured in terms of orders of magnitude. For example, it improves NGA50 by a factor of 5 (HyLight: 351 848; MetaPlatanus: 68 613; Opera-MS: 41 134), indel error rate by a factor of 24 (HyLight: 0.85/100 kbp; MetaPlatanus: 20.56/100 kbp) and mismatch error rate by a factor of 13.7 over the second best approach (HyLight: 23.56/100 kbp; MetaPlatanus: 324.99/100 kbp). It further improves missambled contig rate (MC) by a factor of 8.4 (HyLight: 0.19%; MetaPlatanus: 1.6%).

**Table 1.** Benchmark results for assembly simulated PacBio CLR reads. Indels/100 kbp: average number of insertion or deletion errors per 100,000 aligned bases. Mismatches/100 kbp = average number of mismatch errors per 100,000 aligned bases. Genome Fraction GF reflects how much of each of the strain-specific genomes is covered by contigs. N/100 kbp denotes the average number of uncalled bases (N’s) per 100,000 bases in contigs. MC = fraction of misassembled contigs.

| Assembly | GF(%) | NGA50 | Indels/100 kbp | Mismatches/100 kbp | N/100 kbp | MC(%) |
| --- | --- | --- | --- | --- | --- | --- |
| *3 Salmonella* |  |  |  |  |  |  |
| MetaPlatanus | 72.25 | 68613 | 20.56 | 324.99 | 2.00 | 3.15 |
| Unicycler | 70.92 | - | 109.42 | 1957.53 | 0.00 | 6.88 |
| OPERA-MS | 68.43 | 41134 | 115.57 | 559.08 | 0.18 | 7.69 |
| hybridSPAdes | 46.22 | - | 35.73 | 816.90 | 0.00 | 1.60 |
| HyLight | **96.03** | **351848** | 0.85 | **23.56** | 0.00 | 0.19 |
| Strainberry | - | - | - | - | - | - |
| StrainXpress | 90.99 | 2645 | **0.7** | 59.61 | 0 | **0.07** |
| 20 strains |  |  |  |  |  |  |
| MetaPlatanus | 70.33 | 46882 | 39.11 | 238.79 | 23.21 | 2.15 |
| Unicycler | 68.59 | 46894 | 36.44 | 262.88 | 0.00 | 1.44 |
| OPERA-MS | 66.25 | 40464 | 189.74 | 290.46 | 0.09 | 3.66 |
| hybridSPAdes | 62.01 | 13237 | 11.83 | 333.44 | 0.00 | 0.44 |
| HyLight | 91.76 | **139730** | 3.97 | 59.96 | 0.00 | 0.20 |
| Strainberry | 78.43 | 95338 | 8.48 | **26.26** | 19.60 | 2.30 |
| StrainXpress | **92.74** | 1753 | **1.18** | 44.8 | 0 | **0.04** |
| 100 strains |  |  |  |  |  |  |
| MetaPlatanus | 76.60 | 46937 | 36.80 | 407.42 | 41.46 | 2.05 |
| OPERA-MS | 68.61 | 25223 | 39.31 | 575.26 | 0.53 | 7.90 |
| Unicycler | - | - | - | - | - | - |
| hybridSPAdes | - | - | - | - | - | - |
| HyLight | **93.99** | **163296** | 16.87 | 75.95 | 0.00 | 0.93 |
| Strainberry | 79.25 | 65706 | 210.30 | 115.87 | 68.05 | 4.84 |
| StrainXpress | 91.79 | 2597 | **2.00** | **75.48** | 0 | **0.12** |
| 210 strains |  |  |  |  |  |  |
| OPERA-MS | 71.15 | 43045 | 185.34 | 483.57 | 0.13 | 7.15 |
| MetaPlatanus | - | - | - | - | - | - |
| Unicycler | - | - | - | - | - | - |
| hybridSPAdes | - | - | - | - | - | - |
| HyLight | **90.49** | **128015** | 24.59 | 66.41 | 0.00 | 1.63 |
| Strainberry | 78.56 | 81842 | 418.36 | 102.55 | 41.72 | 4.66 |
| StrainXpress | 89.11 | 1240 | **1.89** | **53.42** | 0 | **0.19** |

##### Strainberry / StrainXpress

See Table 1. Strainberry encountered difficulties in the phasing step due to the high similarity among the three *salmonella* strains (ANI¿99%). As a result, Strainberry could not identify sufficiently many SNPs to separate reads from contigs, as assembled by Metaflye. Consequently, Strainberry was unable to assemble this dataset. StrainXpress achieves Genome Fraction that is superior over all prior approaches, but outperformed by HyLight (90.99%), and achieves excellent results in categories relating to contig quality (errors and misassemblies). However, in terms of contiguity, StrainXpress lags behind all other approaches, by large margins. This is no surprise, of course, since StrainXpress is the only approach that does not make use of long reads, which drastically limits its potential to output longer contigs.

#### 20 bacterial strains

This data set consists of 20 strains from 10 different species, resulting in an average of two strains per species. The average coverage for Illumina (NGS) and PacBio (TGS) reads is 20X and 10X, respectively, again reflecting a TGS low coverage scenario. For further details, please refer to the Methods section.

##### Hybrid Assembly Approaches

See again Table 1. Also on this data set, HyLight outperforms the other four methods across all categories. HyLight achieves a Genome fraction of 91.76%, surpassing the current best method by more than 21% (MetaPlatanus: 70.33%). The NGA50 of HyLight is 139,730, which exceeds the second best NGA50 by a factor of 3 (Unicycler: 46 894). Regarding errors, HyLight’s contigs mark a threefold improvement in terms of indel error rates (HyLight: 3.97/100 kbp; HybridSPAdes: 11.83/100 kbp), and the mismatch error rate is four times lower than the toughest competitor (HyLight: 59.96/100 kbp; MetaPlatanus: 238.79/100 kbp). Finally, the there are only half as many misassembled contigs relative to the second best competing method (HyLight: 0.20%; HybridSPAdes: 0.44%).

##### Strainberry / StrainXpress

As was expected, both Strainberry and StrainXpress are competitive with respect to Genome Fraction. However, while StrainXpress (93.45%) even outperforms HyLight (91.76%), Strainberry achieves 78.43%, which from an overall perspective still is remarkable. Unlike Strainberry’s assembly, whose contiguity is competitive (NGA50 - Strainberry: 95338; HyLight: 139730), StrainXpress’ NGA50 is not competitive (2955). Further, rather unexpectedly, Strainberry’s error rates are largely on par with the low error rates of HyLight and StrainXpress (Indels - HyLight: 5.55/100kbp; Strainberry: 8.48/100kbp; StrainXpress: 1.18/100kbp; Mismatches - HyLight: 58.66/100kbp; Strainberry: 26.26/100kbp; StrainXpress: 44.8/100kbp). While the good error rates of Strainberry can be attributed to the low complexity of the data set, results in the other categories reflect expected outcomes when working with low coverage TGS and/or (medium coverage) NGS data. Here, just as much as on all other data sets, StrainXpress has the lowest misassembled contig rate (0.04% vs. 0.20% by HyLight).

#### 100 bacterial strains

This data set consists of 100 strains from 30 species, at an average coverage of 20X per strain for NGS (Illumina) and the (as usual low) 10X per strain for TGS (PacBio CLR) reads. The data set is designed to reflect a more complex scenario. The idea is to evaluate which of the available approaches potentially become confused if the mix of strains becomes more complex and more diverse.

##### Hybrid Assembly Approaches

See Table 1). In fact, despite a slight increase in terms of errors, HyLight remains unaffected by the elevated complexity and continues to outperform the other four methods. Note first that neither HybridSPAdes nor Unicycler were able to perform the assembly within a month time, so we terminated the corresponding runs (on 32 CPUs and 500 GB RAM) not terminating when the strain number reached 100, short read volume reached 16G, and long read volume reached 10G). As for Genome Fraction, HyLights outperforms the other methods by at least 17%, where GF even exceeds the GF achieved on the low complexity data set (HyLight: 93.99%; MetaPlatanus: 76.6%). The NGA50 exceeds the second best one by 3.5 times (HyLight: 163 296; MetaPlatanus: 46 937). Indel error rates are still lower by a factor of more than 2 (HyLight: 16.87/100 kbp; MetaPlatanus: 36.8/100 kbp) and mismatch error rates are smaller by a factor of more than 5 (HyLight: 75.95/100 kbp; MetaPlatanus: 407.42/100 kbp). Misassembled contig rate is smaller by a factor of more than 2 (HyLight: 0.93; Metaplatanus: 2.05).

Thanks to variations in the average coverage of the TGS data of the 100 strains (average coverage follows a log-normal distribution by the design of CAMI), one can analyze the influence of coverage on the quality of the assemblies of the different strains. See Figure 4 for the corresponding results. In comparison to the other two methods whose runs terminated successfully, HyLight generally achieves greater Genome Fraction across all strains. HyLight’s advantages become particularly noticeable at coverage rates below 20X where HyLight outperforms the other methods by large margins with respect to all categories that refer to strain awareness and error content.

**Figure 4.**
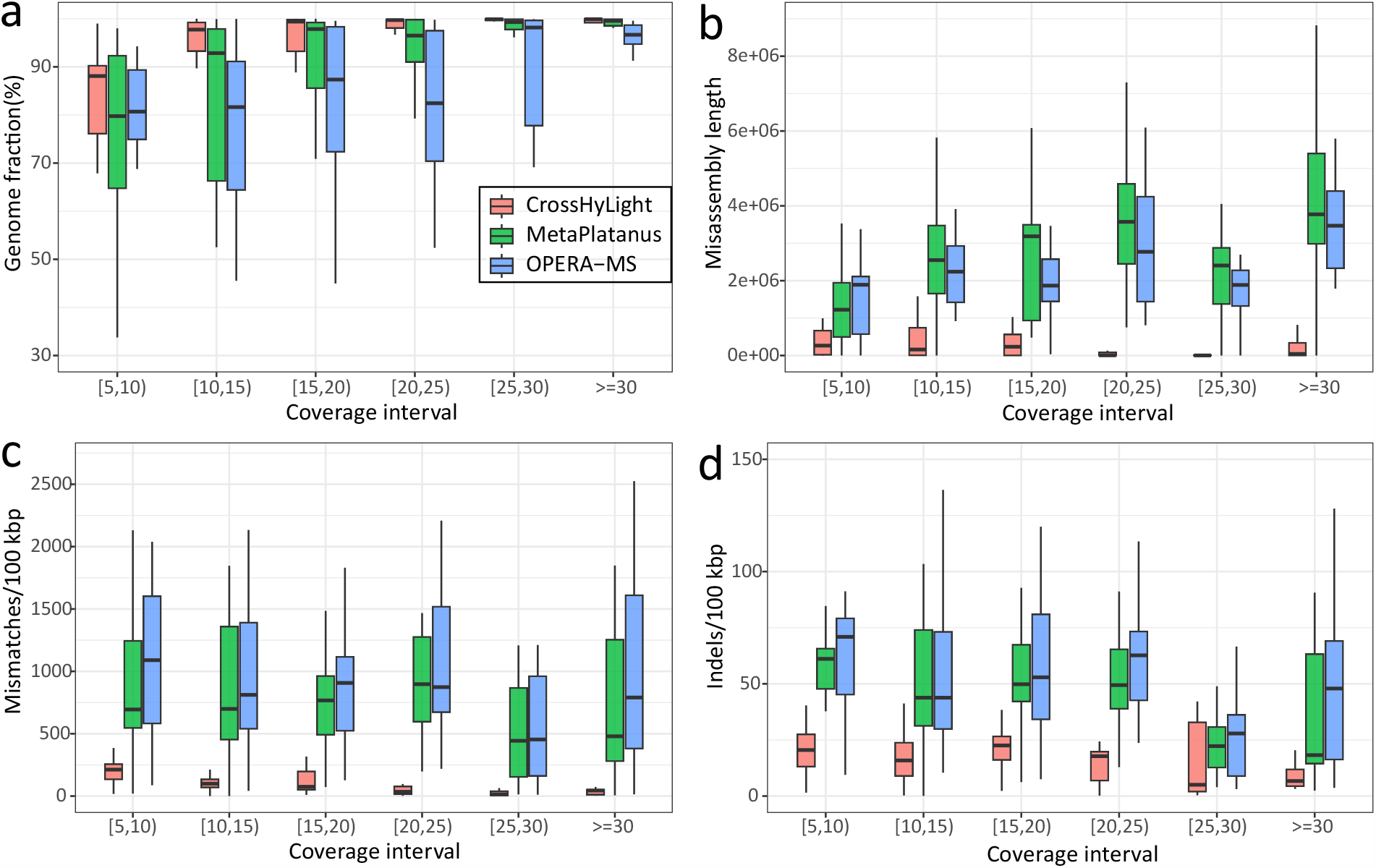
Assemble 100 strains. Among these 100 strains, their average coverages follow a log-normal distribution, resulting in variations in the average coverage of each individual strain. Here, we assess the impact of different coverages on the assembly methods of HyLight, MetaPlatanus, and OPERA-MS. Different colors represent different assembly approach. a: As coverage increases, there is a change in the genome fraction for distinct approaches. b: As coverage increases, there is a variation in misassembly contig length among different assembly methods. c and d: Increase in coverage, changes in mismatch and indel error rate in the assembly results of different approaches.

##### Strainberry / StrainXpress

Again, both approaches prove to be competitive in terms of strain awareness (Genome Fraction - HyLight: 93.86%; Strainberry: 79.25%; StrainXpress: 95.16%). Again StrainXpress even slighly outperforms HyLight, while Strainberry still achieves rmarkable performance rates from a larger perspective. The increased complexity of the data now translates into results in terms of the contiguity and the error rates of the assemblies. Strainberry has considerable disadvantages in terms of error rates (Indels - HyLight: 11.68/100kbp; Strainberry: 210.30/100kbp; StrainXpress: 2.00/100kbp; Mismatches - HyLight: 55.68/100kbp; Strainberry: 115.87/100kbp; StrainXpress: 75.48/100kbp), while StrainXpress has considerable disadvantages in terms of contiguity (NGA50 - HyLight: 163296; Strainberry: 65706; StrainXpress: 2174), whereas HyLight keeps competitive performance both in terms of contiguity and error content.

#### 210 bacterial strains

This data set consists of 210 strains from 100 species as provided by (56), for which reads were simulated using CAMISIM (57). As usual, depth of coverage is 10X for TGS (so low coverage), and 20X for NGS reads. The data set is supposed to reflect a scenario of utmost complexity with respect to numbers of species and their strains.

##### Hybrid Assembly Approaches

See Table1 for results. By and large, HyLight, as well as Opera-MS, as the only methods whose runs terminated within a month time, approximately mirror the results achieved on the data set containing 100 strains. HyLight outperforms Opera-MS by 19% in terms of Genome Fraction (HyLight: 90.95%; Opera-MS: 71.15%), has three times longer contigs (NGA 50: HyLight: 128 015; Opera-MS: 43 045), has more than 3 times lower indel error rates (HyLight: 24.59/100 kbp; Opera-MS: 185,34/100 kbp), 7 times lower mismatch error rates (HyLight: 66.41/100 kbp; Opera-MS: 483.57/100 kbp) and more than 4 times less misassembled contigs (MC: HyLight: 1.63%; Opera-MS: 7.15%).

##### Strainberry / StrainXpress

Results largely repeat the achievements from the data set on 100 strains just discussed, but become even more distinct in terms of the expected advantages and disadvantages of the approaches. Both approaches are competitive in terms of strain awareness (Genome Fraction - HyLight: 90.16%; Strainberry: 78.56%; StrainXpress: ?%), where here, finally, HyLight also clearly outperforms StrainXpress. While Strainberry has drawbacks with respect to error rates (Indels - HyLight: 17.69/100kbp; Strainberry: 418.36/100kbp; StrainXpress: ?; Mismatches - HyLight: 52.78/100kbp; Strainberry: 102.55/100kbp; StrainXpress: ?), StrainXpress considerably trails in terms of contiguity (NGA50 - HyLight: 128015; Strainberry: 81842; StrainXpress: ?).

#### Strain-mixing spike-in datasets

By its design, these data sets can be used to investigate the influence of the coverage of the NGS reads in hybrid assembly. For enabling such experiments, we made use of 10 highly identical *Salmonella* strains, which we spiked into real metagenome samples. While the coverage of spiked-in long reads was fixed to 10X, the coverage of spiked-in NGS reads varied from 5X to 30X, in steps of 5X, resulting in 6 different levels of coverage. These 6 different NGS read sets were spiked into 6 different real metagenome sequencing data sets, amounting to 36 different data sets overall. For each of these 36 data sets, the task is to assemble the genomes of the 10 *Salmonella* strains, in a strain-aware manner.

For the evaluation, note that we do not make use of MetaQuast, because MetaQuast was not able to align the contigs against the reference effectively due to the high identity of many strains, often amounting to average nucleotide identity (ANI) of more than 99%. This implied that indel and mismatch error rates were evaluated as excessive for all methods apart from HyLight (for example, the mismatch error rate was evaluated as 6859.95/100 kbp for Opera-MS, which cannot be correct, see Supplementary Table 1). For fairness reasons, we therefore resorted to using Quast (61) with the same parameters, because we realized that Quast aligned contigs with the reference genomes more accurately. This immediately entailed that the error rate of the competitors dropped substantially (e.g. Opera-MS now at 1753.41/100 kbp). See Supplementary Table 1 for both Quast and MetaQuast evaluated results.

An additional challenge was that MetaPlatanus consistently threw errors when dealing with data sets of only 5X or 10X simulated NGS coverage. Despite reaching out to the authors (via GitHub), the issue could not be resolved. Therefore, we only display results for datasets of simulated NGS coverage 15X and greater.

##### Hybrid Assembly Appraoches

See the Figure5 for results. HyLight outperforms the other methods across different coverages. For example, HyLight achieves an average Genome Fraction that exceeds those of other approaches by at least 28.81% (24.65% ∼26.93%). Note that at coverage 5X, the Genome Fraction of HyLight drops to 77.45%. Genome Fraction for HyLight already increases to 85.03% when increasing coverage to 10X. Increasing coverage further does not lead to any more significant changes. Subsequent increases in the coverage of the most community did not have a more significant changes (Genome Fraction rises to nearly 90% from 15X and onwards).

**Figure 5.**
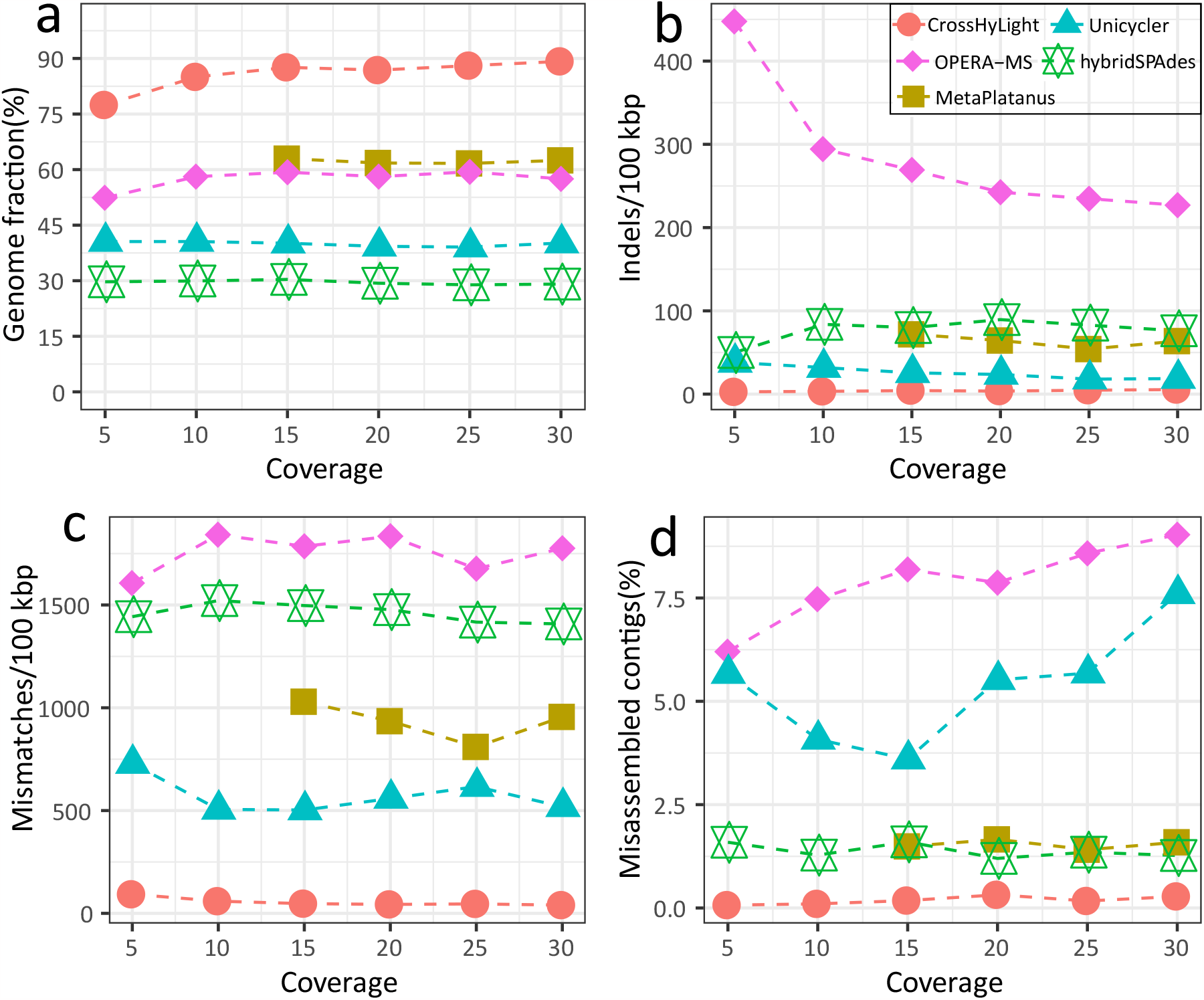
10 *Salmonella* strains spike-in. Simulated reads of 10 *Salmonella* strains were mixed with real sequencing data, followed by assembly for the combined datasets. Subsequently, the quality of assembly results for these spike-in strains in a complex environment was evaluated. During the incremental increase of coverage from 5X to 30X, the variations in genome fraction (a), indel error rate (b), mismatch error rate (c), and misassembly contig rate (d) were presented for these five assembly methods.

Among the prior hybrid assemblers, Unicycler achieves the lowest indel error rate (17.97/100 kbp ∼37.98/100 kbp) where HyLight (2.73 ∼5.47/100 kbp) achieves an error rate of only 17.5% of that of Unicycler (Figure5 b). Improvements of HyLight over prior hybrid assemblers in terms of mismatch errors are even more distinct, decreasing the one of the top competitor by about 90%. Figure5 d finally displays comparison in terms of misassembled contig rate (MC): HyLight’s MC is only 17.5% of that of HybridSPAdes, as the toughest competitor.

In summary, HyLight outperforms the state-of-the-art in hybrid assembly by large margins with respect to the most relevant key assembly metrics, in a scenario that is characterized by near-identical strains embedded into complex real backgrounds.

*Strainberry / StrainXpress* are not evaluated, because the experiment serves the purposes of evaluating particular qualities of hybrid assembly approaches.

### Experiments: Real Data Sets

We conducted further evaluations of all approaches using four real datasets: “Bmock 12 PacBio”, “Bmock 12 ONT”, “NWCs PacBio” and “NWCs ONT”. While TGS coverage of the 4 data sets amounted to 22.11X, 18.1X, 127.2X and 89.01X (in the order of having listed data sets before), NGS coverages reached 275X for Bmock12 and 35.62X for NWCs.

The ‘Bmock12’ dataset consists of 11 strains from 9 species. Due to the low number of strains per species, also strain-unaware approaches are to achieve fairly high Genome Fraction overall. This also means that a sufficiently thorough analysis of the strain awareness of the approaches requires to break down results relative to the strains that make part of the data set. In this vein, one notices that among the species present, only *Marinobacter* and *Halomonas* have more than one strain, see Table 3 for summarizing statistics that refer to “Bmock 12 ONT” (statistics for “Bmock 12 PacBio” are similar, see Supplementary Table S3). While the two strains of *Marinobacter* exhibit 85% average nucleotide identity (ANI), the two *Halomonas* strains have an ANI of 99%. This points out that methods should be evaluated with a view towards their performance on *Halomonas* strains in particular.

The NWC dataset includes 3 species *(Streptococcus thermophilus, Lactobacillus delbrueckii, Lactobacillus helveticus)*, each of which has 2 strains, at ANIs of 99.99%, 99.24%, and 98.03%, respectively. For more detailed information, please see Supplementary Table 4. Despite the limited number of strains and their relatively low complexity, assembly remains challenging due to the high degree of similarity affecting the two strains of a particular species.

### Bmock12 ONT

#### Hybrid Assembly Approaches

See Table 2 for the following results. Due to the reduced level of complexity in terms of the variety of strains, also strain-unaware, species-level metagenome assembly approaches are expected to deliver good performance in reconstructing the individual genomes. Despite the reductions in overall coverage due to the subsampling procedure (inducing a low coverage TGS data scenario), both HybridSPAdes and Unicycler were unable to complete the assembly process within one month runtime.

**Table 2.** Benchmark results for assembly real reads. Indels/100 kbp: average number of insertion or deletion errors per 100,000 aligned bases. Mismatches/100 kbp = average number of mismatch errors per 100,000 aligned bases. Genome Fraction GF reflects how much of each of the strain-specific genomes is covered by contigs. N/100 kbp denotes the average number of uncalled bases (N’s) per 100,000 bases in contigs. MC = fraction of misassembled contigs.

| Assembly | GF(%) | NGA50 | Indels/100kbp | Mismatches/100kbp | N/100kbp | MC(%) |
| --- | --- | --- | --- | --- | --- | --- |
| Bmock12 ONT |  |  |  |  |  |  |
| MetaPlatanus | 95.37 | **789960** | 34.34 | 66.81 | 178.39 | 4.60 |
| OPERA-MS | 94.30 | 207591 | 20.78 | 78.01 | 0.04 | 8.38 |
| hybridSPAdes | - | - | - | - | - | - |
| Unicycler | - | - | - | - | - | - |
| HyLight | **99.77** | 281944 | **1.45** | **3.58** | 0.00 | 3.59 |
| Strainberry | 67.60 | 688598 | 705.34 | 264.57 | 3.57 | 11.66 |
| StrainXpress | 99.04 | 65743 | 34.17 | 41.4 | 19.57 | **0.78** |
| Bmock12 PacBio |  |  |  |  |  |  |
| MetaPlatanus | 94.26 | **809518** | 15.85 | 31.83 | 29.11 | **4.29** |
| OPERA-MS | 93.20 | 179729 | 39.65 | 52.19 | 0.03 | 8.60 |
| hybridSPAdes | - | - | - | - | - | - |
| Unicycler | - | - | - | - | - | - |
| HyLight | **98.57** | 123823 | **5.29** | **19.24** | 0.00 | 7.62 |
| Strainberry | 62.50 | 72377 | 272.41 | 33.48 | 21.30 | 22.17 |
| NWCs ONT |  |  |  |  |  |  |
| MetaPlatanus | 68.54 | 20673 | 260.28 | 248.95 | 2497.22 | 4.03 |
| OPERA-MS | 62.52 | 7656 | 239.54 | 256.18 | 0.38 | 41.54 |
| hybridSPAdes | 56.36 | - | **16.00** | 102.93 | 0.00 | 4.59 |
| Unicycler | 53.78 | 10687 | 117.12 | **68.69** | 0.00 | 34.15 |
| HyLight | **95.35** | 62800 | 30.45 | 174.89 | 0.00 | 9.37 |
| Strainberry | 91.69 | **141570** | 764.14 | 193.84 | 60.26 | 22.94 |
| StrainXpress | 75.47 | 1056 | 25.57 | 399.71 | 0 | **3.38** |
| NWCs PacBio |  |  |  |  |  |  |
| MetaPlatanus | 63.86 | 839 | 116.67 | 143.66 | 802.98 | 4.19 |
| OPERA-MS | 56.49 | - | 446.73 | 315.74 | 0.41 | 26.57 |
| hybridSPAdes | 54.83 | - | **22.53** | 124.98 | 0.00 | **3.57** |
| Unicycler | 46.96 | - | 36.14 | **57.18** | 0.00 | 11.06 |
| HyLight | **78.94** | **22388** | 84.42 | 219.74 | 0.00 | 4.27 |
| Strainberry | 43.03 | - | 246.94 | 111.47 | 0.65 | 22.22 |

**Table 3.** The genome fraction of each individual strain in the Bmock12 data (Illumina and ONT). Present the impact of different sequencing coverage in different assembly methods.

| Assembly | Coverage (Illumina) | Coverage (ONT) | HyLight | MetaPlatanus | OPERA-MS |
| --- | --- | --- | --- | --- | --- |
| **Halomonas sp.HL-4** | **507.08** | **46.03** | **99.63** | **93.05** | **90.70** |
| **Halomonas sp.HL-93** | **579.87** | **41.23** | **99.68** | **96.75** | **93.90** |
| **Marinobacter sp.LV10R510-8** | **447.83** | **18.19** | **100.00** | **99.97** | **99.80** |
| **Marinobacter sp.LV10MA510-1** | **135.05** | **6.95** | **99.94** | **99.85** | **98.95** |
| Muricauda sp.ES.050 | 618.76 | 53.87 | 100.00 | 99.92 | 99.82 |
| Psychrobacter sp.LV10R520-6 | 425.47 | 31.82 | 100.00 | 99.34 | 98.46 |
| Cohaesibacter sp.ES.047 | 170.59 | 11.78 | 99.78 | 99.48 | 98.57 |
| Thioclava sp.ES.032 | 78.32 | 5.54 | 99.64 | 99.65 | 99.40 |
| Propionibacteriaceae bacterium | 31.90 | 2.39 | 99.99 | 99.99 | 99.97 |
| Micromonospora echinofusca | 18.19 | 2.44 | 99.58 | 99.50 | 99.29 |
| Micromonospora echinaurantiaca | 14.91 | 1.80 | 99.64 | 99.67 | 99.11 |

HyLight outperforms both Opera-MS and MetaPlatanus, which are the two hybrid assembly approaches whose runs terminate in acceptable time when examining the relevant criteria and putting them into mutual perspective. Genome Fraction of HyLight is 4.4% higher than that of the second-ranked MetaPlatanus (99.77% vs. 95.37%). MetaPlatanus achieves the greatest NGA50 (789,960 vs. 281,944), which, however, can be explained by the unusually large number of Ns in its contigs, whose primary purpose is to link and extend contigs by force without that read evidence for the missing sequence context of contig links can be provided. HyLight improves by more than one order of magnitude over the other methods in terms of indel and mismatch errors. The indel error rate of HyLight is only 6.9% of that of the second-ranked OPERA-MS (1.43 vs. 20.78 per 100 kbp), and the mismatch error rate of HyLight is only 5.4% of that of the second-ranked MetaPlatanus (3.58 vs. 66.81 per 100 kbp).

We further examined the assembly status for each strain making part of the mock community individually. Genome Fraction for the individual strains is displayed in Table3. One immediately realizes that all approaches reconstruct at least (about) 99% of the strain-specific sequence for all but the two *Halomonas* strains whose ANI comes at 99%. On *Halomonas sp*.*HL-4* in particular, HyLight achieves a Genome Fraction that is greater by 7.58% than that of MetaPlatanus and 8.93% than that of Opera-MS (HyLight: 99.36; MetaPlatanus: 93.05; Opera-MS: 90.7). This lets one conclude that HyLight is the only hybrid metagenome assembly approach that operates in a strain-aware manner even when the ANI between strains is as great as 99%. As an additional insight gained from the analysis of the quality of the assemblies that is stratified relative to the individual strains, one realizes that all hybrid assembly approaches are able to reconstruct about 99% of the strain specific sequence even when TGS sequencing coverage is as low as 1.8X and also as low as 14.91X for the respective NGS coverage for a particular strain (here: *Micromonospora echinaurantiaca*). This documents the general value of hybrid metagenome assembly with respect to its favorable behavior in terms of sequencing (in particular TGS) coverage demands.

#### Strainberry / StrainXpress

While StrainXpress achieves competitive performance, Strainberry somewhat unexpectedly, does not (Genome Fraction - HyLight: 99.77%; Strainberry: 67.60%; StrainXpress: 99.04%). While Strainberry has (here: drastic) drawbacks with respect to error rates (Indels - HyLight: 1.45/100kbp; Strainberry: 705.34/100kbp; StrainXpress: 34.17/100kbp; Mismatches - HyLight: 3.58/100kbp; Strainberry: 264.57/100kbp; StrainXpress: 41.4/100kbp), StrainXpress considerably trails in terms of contiguity (NGA50 - HyLight: 281944; Strainberry: 688598; StrainXpress: 65743), while Strainberry even outperforms HyLight.

##### Bmock12 PacBio

Note that NGS reads used here agree with those from “Bmock12 ONT”, whereas the TGS reads now stem from PacBio CLR sequencing platforms. This means in particular that the results of StrainXpress (see further below agree with those achieved on “Bmock12 ONT”.

###### Hybrid Assembly Approaches

See Table 2 for results. In an overall summary, results here mirror results achieved on “Bmock12 ONT”, where advantages of HyLight look less drastic: HyLight maintains a Genome Fraction that exceeds that of the other approaches by more than 4%, and although less distinct, still exhibits considerably lower error rates. The NGA50 of MetaPlatanus exceeds that of HyLight considerably, again put into context by the large number of N’s in the MetaPlatanus contigs. Misassembled contig rate of HyLight is again roughly on a par with that of MetaPlatanus, where now MetaPlatanus has slight advantages (but see below for a more fine-grained analysis that provides explanations). Although results look more favorable for MetaPlatanus from a greater persepctive, breaking down results by strain (see Supplementary Table 3) reveals that MetaPlatanus substantially struggles in reconstructing one of the *Halomonas* strains: while achieving 93.05 Genome Fraction for *Halomonas sp*.*HL-4* on ONT data, MetaPlatanus only achieves 82.19 Genome Fraction on PacBio data, despite even having small advantages over HyLight on the other *Halomonas* strain. Analogous trends become evident for Opera-MS. With respect to the misassembly contig rate, a more detailed analysis further demonstrates that the MC of the raw long reads (i.e. evaluating raw long reads as contigs in their own right) for “Bmock12 ONT” and “Bmock12 PacBio” comes out at 2.33% and 7.55%, see Supplementary Table 2. The MC of the raw long reads are introduced by chimera reads. Comparing the MC of the raw long reads with the MC of HyLight points out that HyLight reproduces MC rates of the raw long reads. The most plausible explanation for this is the fact that HyLight uses overlap graphs, which cannot identify chimera reads, while “short-read-first” approaches, thanks to employing DBG’s, can identify artificial links as mistaken. While there is good hope that overlap graph based approaches are able to identify chimera reads, too, we leave such improvements as promising future work at this point.

###### Stainberry / StrainXpress

StrainXpress reproduces its results because making use of only the NGS portion of the data, which agrees with that from “Bmock12 ONT”. Again, Strainberry somewhat unexpectedly, does not achieve competitive performance in terms of strain awareness (Genome Fraction - HyLight: 98.57%; Strainberry: 62.50%). Again, Strainberry has considerable drawbacks with respect to in particular indel error rates (Indels - HyLight: 5.92/100kbp; Strainberry: 272.41/100kbp; StrainXpress: ?; Mismatches - HyLight: 19.24/100kbp; Strainberry: 33.48/100kbp; StrainXpress: ?), while the contiguity of Strainberry’s assembly here is outperformed by that of HyLight (NGA50 - HyLight: 123823; Strainberry: 72377).

#### NWC ONT

##### Hybrid Assembly Approaches

Due to the presence of a higher number of highly similar strains in NWC compared to Bmock12, the advantage of HyLight becomes more pronounced. HyLight drastically outperforms the other hybrid assembly methods, both in terms of Genome Fraction (i.e. strain awareness; HyLight: 95.35%; MetaPlatanus, as second best: 68.54%) and NGA50 (i.e. contiguity; HyLight: 62800; MetaPlatanus, as second best: 20673). Further, although not outperforming the other approaches, HyLight achieves decent indel and mismatch error rates in comparison with the other approaches, ranking second and third, respectively, and, just like most other approaches, reducing the error content of the raw long reads by more than two orders of magnitude. Again, the MC, although not bad, is slightly worse than that of MetaPlatanus and HybridSPAdes, again reflecting that HyLight adopts issues introduced by chimera reads, for which there is good hope that this can be successfully addressed in future work.

##### Strainberry / StrainXpress

All approaches achieve competitive performance relative to strain awareness, in case of StrainXpress at least from the perspective of comparing it with strain unaware approaches, as was expected (Genome Fraction - HyLight: 95.35%; Strainberry: 91.69%; StrainXpress: 75.29%). Strainberry has (drastic) drawbacks with respect to indel error rates (Indels - HyLight: 30.45/100kbp; Strainberry: 764.14/100kbp; StrainXpress: ?; Mismatches - HyLight: 174.89/100kbp; Strainberry: 193.84/100kbp; StrainXpress: ?), StrainXpress considerably trails in terms of contiguity (NGA50 - HyLight: 62800; Strainberry: 141570; StrainXpress: 636), with Strainberry taking the lead.

#### NWC PacBio

Unlike NWC ONT, this data set is affected by two strains whose long read coverage is extremely low (1.45X and 2.47X, resepectively, see Supplementary Table 6). The reconstruction of those two strains presents a particular challenge, which points out that this data set is a particularly challengin one overall. Therefore, all methods produced assembly results that were inferior to those achieved on NWC ONT.

##### Hybrid Assembly Approaches

Notwithstanding the level of difficulty of the data set overall, results virtually reproduce the ones achieved on NWC ONT: HyLight drastically outperforms the other approaches both in terms of Genome Fraction (HyLight: 78.94%; MetaPlatanus: 63.86% as second best) and NGA50 (HyLight: 22388; MetaPlatanus: 839 as second best). Error rates are roughly on a par with those of the other approaches, where everyone achieves sufficiently decent results. MC is slightly lower than those of the two best approaches, but considerably better than those of the other two approaches; again, presumably, chimera reads imply that HyLight reproduces the MC rates of the raw reads.

##### Strainberry / StrainXpress

Just as for Bmock12, StrainXpress reproduces its results because making use of only the NGS portion of the NWC data. Again, Strainberry’s performance in terms of strain awareness drops (here: quite substantially), which is somewhat unexpected, and may be due to reduced quality of the TGS portion of the data here—note that even HyLight somewhat struggles (Genome Fraction - HyLight: 78.94%; Strainberry: 43.03%). In comparison to “NWC ONT”, Strainberry’s error rates are improved (Indels - HyLight: 84.42/100kbp; Strainberry: 246.94/100kbp; StrainXpress: ?; Mismatches - HyLight: 219.74/100kbp; Strainberry: 111.47/100kbp; StrainXpress: ?). In terms of contiguity, HyLight clearly outperforms Strainberry (NGA50 - HyLight: 22388; Strainberry: - because its contigs do only align with less than 50% of the true genomes).

### Runtime and memory usage evaluation

We evaluated the performance of runtime and peak memory of all methods on the data set containing the 3 *Salmonella* strains, on a x86 64 GNU/Linux machine with 48 CPUs. The data volume of the NGS (Illumina) reads amounted to 573 MB and the volume of the Pacbio CLR reads amounted to 281 MB. Supplementary Table S2 reports CPU times and peak memory usages of the different hybrid assembly methods. without any doubt, OPERA-MS is the fastest tool: it only takes 2.09 hours and 1.23 GB memory. The runtime of hybridSPAdes, HyLight, and MetaPlatanus is roughly on a par, requiring approximately 5.53, 7.01, and 6.93 hours, respectively. However, in terms of peak memory usage, both hybridSPAdes and HyLight demonstrate significantly lower usage compared to MetaPlatanus, with values of 3.85, 15.99, and 69.26, respectively. Unicycler, on the other hand, requires the longest runtime (53.71 hours).

## Discussion

Despite exhibiting a high degree of similarity, different strains of the same bacterial species can vary significantly in terms of phenotype, such as drug resistance or pathogenicity. This explains why it is important in biomedical and clinical research to identify putative pathogens at the level of strains, and not only at the level of species, as the desirable degree of texonomic resolution. So, when analyzing an environmental mix of genomes, the current driving challenge is to assemble the individual genomes at strain resolution, *De novo* assembly is crucial to avoid reference induced biases that would favor the detection of more common strains over those not yet fully investigated or even entirely unknown.

In this paper, we have presented HyLight, as a novel approach to push the limits of *strain-aware metagenome assembly* in a substantial manner. HyLight is based on de novo *hybrid assembly*, characterized by integrating both long, third-generation sequencing reads and short, next-generation sequencing reads during the assembly process. Two good reasons support the principled superiority of hybrid assembly.

### The first reason is the superiority of the assemblies themselves

HyLight improves the quality of strain-level metagenome assemblies in comparison with the state of the art all in terms of strain awareness, contiguity, and accuracy (as measured by error and misassembly content), often by large margins. As anticipated, HyLight achieves its most pronounced advantages when strains are very similar and/or when strains are subject to low (TGS) read coverage. HyLight stands out as the sole method that consistently reconstructs at least 90% of the strain-specific sequence content, with second best approaches achieving only approximately 70% on average. Another particular advantage of HyLight is the low error content in terms of insertions and deletions, as the predominant type of errors that affects TGS reads. Beyond clearly outperforming competitors in terms of indel error content, HyLight shares the very favorable mismatch error rates that the other approaches had been able to achieve already, which documents the beneficial complementarity of the TGS reads on the one hand and NGS reads on the other hand.

### The second reason are the substantial savings in terms of costs and resources

From the point of view of the global assembly/sequencing community, hybrid assembly opens up a range of opportunities for laboratories that operate on less generous budgets. That is, in other words, it opens up opportunities for the vast majority of laboratories worldwide. The principled explanation for this is the desirable degree of complementarity of the two types of data involved: while TGS reads are long but inaccurate, and in particular predominantly affected by indel errors, NGS reads are short and accurate where the (anyway little) errors come in form of presumable point mutations. This drastically reduces the need for generating larger data volumes. In line with earlier hybrid assembly approaches, HyLight again demonstrates that superior results are achieved already with little and cheap data. Importantly, nearly every sequencing laboratory today counts third-generation-sequencing of the cheaper type (that is more erroneous, but still generating by far the longest reads thanks to the simple sequencing protocols guiding the generation process), and next-generation-sequencing platforms among its standard equipment.

This is in obvious contrast to the latest and most advanced sequencing platforms such as PacBio HiFi or ONT Q30+ type of reads. It seems to be a fair assessment to consider these most advanced types of reads as convenience that remains unaffordable to many. Also, still—see the first reason—the quality of hybrid assemblies exceeds the quality of assemblies computed by such high-convenience protocols. The reason for the latter is the fact that TGS reads of the primary type are just still substantially (by factors of up to 2-4 times) longer. Also, the polishing of TGS reads using indel-error-free NGS reads appears to be advantageous over the refined sequencing protocols that characterize the more convenient technologies.

Here, we have presented an approach that leverages the complementarity of the two types of data sources, ultra-long, though highly erroneous TGS reads on the one hand, and short NGS reads that still constitute the cleanest source of sequencing data on the other hand. Confirming the theoretical arguments, we have demonstrated to outperform the state of the art on low data volumes in particular, often by large margins. So, as intended, we have indeed made it possible to compute strain aware metagenome assemblies for laboratories running on less generous budgets, and we have also substantially pushed the limits in strain aware metagenome assembly in general.

Key to success has been the observation that a considerable remodeling of hybrid assembly protocols was necessary. It had been agreed to classify hybrid assembly approaches into “short-read-first” and “long-read-first” approaches, which either use short reads or, respectively, long reads as basis for their assemblies, while using the other type of data for auxiliary purposes, either to lay out and scaffold the shorter assemblies, or, respectively, to remove errors from the already longer assemblies.

HyLight does not make use of either such conventional strategy. Instead, HyLight is rooted in a “cross hybrid” strategy: it assembles long reads using short reads as auxiliary source of data, and vice versa assembles short reads assisted by long read information. This “mutual support strategy” enables one to resolve repetitive elements and fill gaps in the long read assemblies, which comes in addition to the conventional advantages of (”unilateral support”) hybrid assemblies. We recall that long TGS reads are supposed to have low coverage to save expenses, which provided particular motivation for using the short read data for repeat resolution and gap filling, and not only polishing errors.

To make all of this possible, HyLight employs overlap graphs as the driving underlying data structure. HyLight realizes that the presence of long reads renders usage of de Bruijn graphs obsolete. While this is understood for long read assemblies—as overlap graphs have regained a prominent role when processing long reads—this may be somewhat surprising when considering short reads. As had already been pointed out in prior work, using overlap graphs for assembling short reads in contexts that aim at strain awareness aids in the correct line up of strain specific variation within strain specific genomes.

A particular technical, novel feature of HyLight is to incorporate a filtering step that identifies mistaken (strain-unaware) overlaps and removes them from the graphs. The filtering step prevents the incorrect compression of strain-specific variation into contigs that mistakenly connect sequence from different strains. This particular feature has been key to correctly discerning genomic differences at the level of strains within microbial communities.

In conclusion, we have introduced HyLight, as a novel de novo hybrid metagenome assembly approach that reconstructs the genomes of the individual members of microbial communities at strain level, which is a novelty. An additional feature of potential practical interest is its economic behavior in terms of costs and times, and the ubiquitous availability of the data HyLight relies on. In that vein, HyLight has the potential to deliver strain aware metagenome assemblies to many, likely the large majority of laboratories that had not been able to reconstruct genomes of microbial communities at strain level so far.

Despite the various advantages that we have been able to demonstrate, there is still room for improvement. So far, for example, we have not yet addressed the existence of long chimera reads, which introduce a detectable amount of misassemblies and confound the correct identification of strain specific genomes to a small, but non-negligible degree. Last but not least, even though necessary because key to success, overlap graphs are “heavy” data structures. In future work, we will focus on the design of “lightweight” overlap graphs that, although agreeing on minor amounts of inaccuracies, still achieve superior results while, however, making further substantial savings in terms of runtime and peak memory requirements.

## Methods

### Quality control

Before assembling reads with HyLight, we performed quality control on the sequencing reads using *fastp* (version 0.20.1) (62). This multifunctional FASTQ data preprocessing toolkit ensures the quality of the data by providing major functions including quality control, adapter detection, base correction, and read filtering. In the raw reads, bases with Phred scores less than 20 at the 5’ or 3’ ends, as well as adapters, were trimmed. After trimming, only reads longer than 70 bp were retained. Moreover, in the overlapped regions of paired-end reads, *fastp* corrected mismatched bases only when a high-quality base was paired with a low-quality base.

### Workflow: Detailed Description of 3 modules

#### Module 1: Strain-Aware Assembly of Long Reads

##### Hybrid correct long reads and establish OG

As long-read technologies such as CLR or ONT often contain a significant amount of sequencing errors, which can affect downstream SNP identification and assembly, the first step in our pipeline is to use FMLRC2 (50) to perform error correction on long reads using high-quality short reads. After obtaining high-quality long reads, we use minimap2 to align them to each other and construct an OG. In this OG, each read is represented as a node, and the overlapping regions between them are represented as edges.

##### Untangle wrong overlaps with SNPs

Once a long read OG is established, further optimization is necessary to obtain a strain-resolved OG,. In particular, the OG established contains a considerable amount of mistaken overlaps. The reasons for such mistaken overlaps to show are the high similarity between different strains, which leads to alignment algorithms aligning regions that look similar, but stem from different stains with each other. Therefore, assembling genomes using the still raw OG is prone to accumulating misassemblies and losses of strain-specific variations.

To enable HyLight to assemble strain-resolved contigs, it is therefore crucial to untangle such mistaken overlaps (a.k.a. removing the corresponding edges in the OG). To discover and untangle such overlaps, we utilize SNP information as per the steps displayed in Figure 2.

First, for each read, we keep track of the mismatches and indels in comparison with the reads that aligned with it. Corresponding details can be immediately obtained from examining the CIGAR strings output by Minimap2. In other words, we leverage the mismatch information in combination with counts of reads that support the mismatch characters (nucleotides) to determine SNP sites.

There can be two cases: 1) A base in a read is different from the consensus base in all other reads, see ‘T’ at P1:X and ‘C’ at P2:X in Figure 2. If less than 3 reads support such a base, we consider it a sequencing error. We do not break overlaps between reads due to this case. 2) There are two bases showing, and a sufficient amount of reads (*≥* 3) that support each of them. See Figure 2: this applies for both ‘A’ and ‘C’ at P1:X, ‘T’ and ‘A’ at P2:X, and ‘C’ and ‘T’ at P3:X. In any such case, we break overlaps that are affected by this scenario, in other words, we remove the corresponding edges from the OG.

In case of both 1) and 2) applying for pairs of aligned reads (for example for the read that contains ‘T’ at P1:X in Figure 2), we determine the identity of a read based on the SNP information referring to other positions, where bases of a read were not evaluated as sequencing errors (here P2:X and P3:X).

Note that Figure 2 only refers to the case of reads from two different strains alone. In reality, scenarios encompass more than just two strains, however. Nevertheless, the rules that we have described can be straightforwardly extended to scenarios reflecting the presence of more than 2 strains: we distinguish between sequencing errors (supported by at most 2 reads) and strain-specific variation (supported by at least 3 reads) in the exact same way.

The encompassing evaluation of the SNP information obtained from studying the alignments of the overlapping part of reads eventually leads to unambiguous resolution of strain-aware overlap information, which leads to establishing a strain-aware OG as a result.

##### Assemble long reads by strain aware OG and correct contigs

After obtaining the strain-resolved graph, we can use Miniasm (54) to assemble contigs. Note that standard usage of Miniasm consists in providing an OG as immediately computed by Minimap2. The particular, novel turn applied here is to break overlaps that connect reads from different strains *before* providing the OG as input to Miniasm. Despite prior distinguishing between errors and true variation during mistaken overlap removal, long reads often still contain numerous sequencing errors, which necessitates further correction of the contigs assembled by Miniasm using the strain-resolved OG we provided as input.

To do that, we align the long reads with the contigs and, subsequently, following the strategy described earlier, resolve erroneous overlaps between reads and contigs. After untangling the erroneous overlaps, we apply Racon (55) to correct the contigs further. As per its protocol, Racon cuts the mapped reads into 500bp windows and rapidly corrects contigs by way of a de Bruijn Graph (DBG) based procedure. It is important to note that both Miniasm and Racon originally lacked the ability to perform strain-aware assembly and correction. If the original, raw OG had been provided as input, both Miniasm and Racon would lose strain-specific variations during correction (Racon) and assembly (Miniasm). The outcome would be merged contigs affected by considerably more misassemblies. Since we provide a strain-resolved OG that has retained strain-specific variants correctly, HyLight is not affected by this issue.

#### Module 2: Strain-Aware Assembly of Short Reads

##### Pick up and strain-aware assembly of unmapped short reads

The low depth of long reads implies that still some sequencing errors have remained undetected. The low depth induces further that certain genomic regions or (even entire) strains cannot be reconstructed using long reads alone. To this end, we use short reads that remained unaligned with any of the long reads, to polish long read based contigs further, and fill gaps between the contigs. The specific steps are shown in Figure 3. First, we use Minimap2 to align the short reads to the contigs generated from long reads, creating an OG. Then, based on the strategy mentioned above, we use SNPs to untangle the wrongly linked overlaps between two strains and obtain a strain-resolved OG. Reads that are not in this OG belong to regions or strains that have not yet been reconstructed. We then use the previously published method StrainXpress (12) to cluster these remaining short reads and perform strain aware assembly using the OG’s emerging from the clustered short reads.

#### Module 3: Global assembly

After having computed long read and short read contigs, we aim to extend them further by constructing a global contig graph through contig-to-contig alignment. Each vertex corresponds to a contig, and edges correspond to high-quality overlaps: overlaps are supposed to be longer than 100 bp showing at least 0.99 similarity, following guidance provided by previous works (48; 63)). To reduce complexity, all transitive edges are removed, and contigs are joined into “branch-less” sequences. To identify branches, we evaluate short reads that match the corresponding contigs. After removing branches, HyLight updates the graph and extends contigs further. This process is iterated until no further branches are observed, resulting in the “master contigs” that establish HyLight’s final output, available for downstream functional analysis.

### Synthetic data sets

To compare the performance of different approaches, we utilized CAMISIM (57) (version 0.0.6) to produce four simulated hybrid sequencing datasets consisting of Illumina MiSeq and PacBio CLR. These datasets included 3 *Salmonella* strains, 20 bacterial strains (10 species), 100 bacterial strains (30 species), and 210 bacterial strains (100 species), respectively. CAMISIM is a widely used metagenome simulator capable of modeling second and third-generation sequencing data with varying abundances and multi-sample time series based on real strain-level diversity.

The length of the simulated Illumina MiSeq reads is 2X250 bp, at an insert size of 450 bp. The N50 of PacBio CLR reads is 10kbp at an average sequencing error rate of 10%. As per the principles of CAMISIM, the abundance of different strains is uneven, as sampled from a log-normal distribution. The average coverage of both Illumina MiSeq and PacBio CLR data for the four simulated communities is 20X and 10X respectively. The genomes of the 3 *Salmonella* strains were obtained from earlier work (64). The genomes for the 20 bacterial strains, the 100 bacterial strains and the 210 bacterial strains communities were downloaded from an earlier study (56). For details with respect to Genome ID’s and their average nucleotide identity (ANI), please see Supplementary Table S1.

Additionally, to assess the impact of long read coverage, we generated six sequencing datasets by combining simulated and real data, which represents a typical simulation scenario known as “spike-in” data. This approach allows us to evaluate how methods assemble the simulated (”spiked-in”) data, for which the ground truth is known, within a realistic context (although ground truth is lacking for the real data, hence the need to incorporate simulated reads). Specifically, we incorporated simulated reads from ten well-known *Salmonella* strains downloaded from (64) into six distinct real gut metagenome sequencing datasets. These datasets were obtained from experiments aiming at identifying functional characteristics of low-abundance and uncultured species in the human gut (65) (project number: PRJNA602101).

To simulate reads from the *Salmonella* strains, we utilized the CAMISIM simulator while closely matching the properties of the real sequencing data, ensuring optimal comparability. The synthesized short reads were set to a length of 2X150 bp. To account for the influence of read coverage, the coverage of synthesized long reads for the spiked-in strains varied from 5X to 30X in increments of 5X across the six real data sets. Each of the six spiked-in real data sets represented a specific coverage level for short reads. On the other hand, the coverage of the simulated NGS reads remained fixed at 20X across the six real data sets.

Considering the substantial number of reads, we randomly extracted 8,166,722 NGS reads and 181,092 TGS reads from each real hybrid sequencing data set for a less computationally intensive evaluation. These extracted reads were further processed for analysis. For more details regarding the ten *Salmonella* Genome ID’s and SRA identifiers, please refer to Supplementary Table S1, “spike-in *Salmonella*”.

### Real data sets

We considered two microbial communities for which both TGS and NGS data were available in our experiments:

**Bmock12** is a mock community comprising 12 bacterial strains from 10 different species (66). The mock community was sequenced using all ONT MinION, PacBio and Illumina sequencing platforms. The corresponding data sets were obtained from SRA (illumina SRR8073716, ONT SRR8351023, PacBio SRR8073714). The N50 read length for ONT and PacBio reads is 22,772 and 8,701, respectively. The Illumina reads have a read length of 2X150 bp, with an average insert size of 302.7 bp. It is worth noting that the number of reads mapped to *Micromonospora coxensis*, one of the 12 strains, was negligible (66), so we effectively dealt with only 11 bacterial strains. The average coverage for these 11 strains ranges from 74.56X to 3,093.79X, with a median of 1,376.35X. For the sake of a less runtime-intense evaluation in the light of the large amount of duplicates among the reads, we randomly extracted 20% of the reads, and further processed only these. Lastly, it is important to address the challenges posed by this data set, which involve assembling the long reads of two species whose strains exhibit high average nucleotide identities (ANI). Specifically, this applies to the *Marinobacter* species and the *Halomonas* species, as they contain pairs of strains characterized by ANI values of 85% and 99%, respectively.

#### NWCs

The second real microbial community selected for analysis originates from natural whey starter cultures (NWCs) (67). The metagenome samples of the NWCs were sequenced using Illumina MiSeq, generating reads with a length of 2x300 bp. Additionally, PacBio and ONT sequencing platforms were employed. We acquired the sequence data sets from SRA (illumina SRR7585899 and SRR7589561, ONT SRR7585900, PacBio SRR7589560). The N50 read lengths for PacBio and ONT TGS data are quite similar, measuring 11,895 and 9,562, respectively. In a previous study, complete genomes of six bacterial strains from three species were obtained (67). The GenBank accession numbers for these six genomes are CP029252.1, CP031021.1, CP031024.1, CP031025.1, CP029252.1, and CP031021.1. We utilized these reference genomes as the ground truth for evaluating the accuracy of assembly results for distinct approaches.

In the NWCs datasets, we performed a removal of low-quality bases (*>*Q20), the NGS (Illumina) reads retained an unusually high error rate (indel error rate: 19.58/100 kbp; mismatch error rate: 883.28/100 kbp.). This does not reflect standard scenarios, and can have a considerable impact on the accuracy of the assembly. To restore a standard scenario, we corrected the NGS reads prior to hybrid assembly, by using bfc (68), an approved error corrector applicable for Illumina short reads.

### QUAST Evaluation Criteria

During the evaluation process, we took into account all pertinent categories provided by MetaQUAST V5.1.0rc1 (60), a widely recognized tool for assessing assembly quality. Following established guidelines, we incorporated the flags –ambiguity-usage all and –ambiguity-score 0.9999 specifically for the evaluation of metagenomic data. The remaining parameters were kept at their default values. In the subsequent sections, we will provide concise definitions of the metrics under consideration. For more comprehensive explanations, please refer to http://quast.sourceforge.net/docs/manual.html.

`Indels/100 kbp.` The sequence derived from raw PacBio CLR and ONT reads is susceptible to a significant presence of indel errors. In this context, “indels per 100 kbp” refers to the average count of insertion or deletion errors per 100,000 aligned bases within the contigs.

`Mismatches/100 kbp.` This metric represents the average count of mismatch errors per 100,000 aligned bases in the examined contigs.

`N/100 kbp.` This indicates the average count of uncalled bases (N’s) per 100,000 bases in the evaluated contigs.

`Genome Fraction (GF).` GF represents the proportion of aligned bases in the reference genome to which the contigs are aligned. Essentially, GF indicates the extent to which the evaluated contigs cover each of the strain-specific genomes.

`Misassembled contigs (MC).` A contig is categorized as a *misassembled contig* if it contains one or more misassembly events. A misassembly event is identified when a contig aligns to the correct sequence but exhibits a gap larger than 1 kbp, an overlap exceeding 1 kbp with a different strand, or even with a distinct strain. The percentage of misassembled contig, relative to the total number of evaluated contig, is reported as “misassembled reads.”

`NGA50.` NGA50 is defined as the longest contig length, where all contig of that length or longer align to at least 50% of the true sequence. In other words, NGA50 represents the maximum contig length that provides coverage to at least half of the true sequence through their alignments.

## Code Availability

The source code of HyLight is GPL-3.0 licensed, and publicly available at https://github.com/HaploKit/HyLight.

## Acknowledgements

Not applicable.

## Author Contributions

XK and AS developed the method. XK, XL, and AS wrote the manuscript. XK, WZ and XL conducted the data analysis. XK implemented the software. All authors read and approved the final version of the manuscript.

## Competing Interests

The authors declare that they have no competing interests.

## Supplementary

Supplementary.tex — This contains all supplementary materials referenced in the main text

## Notes

### Competing Interest Statement

The authors have declared no competing interest.

