## Supplementary Material for "HyLight: Strain aware assembly of low coverage metagenomes"

### HyLight: Strain aware assembly of low coverage metagenomes
#### Supplementary Information

Xiongbin Kang[1,2], Wenhai Zhang[1], Xiao Luo[1,*], Alexander Schönhuth[2,*]

[1] College of Biology, Hunan University, Changsha, China.
[2] Genome Data Science, Faculty of Technology, Bielefeld University, Bielefeld, Germany.

*To whom correspondence should be addressed.

#### Supplementary Tables and Figures

**Supplementary Table 1.** The GenBank number of reference genomes. Samples information and assembly results of real gut metagenome sequencing data.
| Supplementary Table 1.xlsx |

**Supplementary Table 2.** The quality of four real long reads datasets was assessed using Quast.

| Raw reads | MC(%) | Indels/100kbp | Mismatches/100kbp | GF(%) | N50 |
|---|---|---|---|---|---|
| Bmock12 ONT | 2.33 | 4001.58 | 2659.11 | 93.63 | 22772 |
| Bmock12 PacBio | 7.55 | 7790.78 | 1851.43 | 87.98 | 8701 |
| NWCs ONT | 7.18 | 7991.02 | 5476.30 | 100.00 | 9918 |
| NWCs PacBio | 7.54 | 11479.15 | 5270.85 | 81.61 | 11725 |

**Supplementary Table 3.** The genome fraction of each individual strain in the Bmock12 data (Illumina and PacBio). Present the impact of different sequencing coverage in distinct assembly methods.

| Assembly | Coverage (Illumina) | Coverage (PacBio) | HyLight | MetaPlatanus | OPERA-MS |
|---|---|---|---|---|---|
| **Halomonas sp.HL-4** | **507.08** | **40.01** | **92.73** | **82.19** | **62.44** |
| **Halomonas sp.HL-93** | **579.87** | **45.99** | **95.57** | **98.54** | **94.92** |
| **Marinobacter sp.LV10R510-8** | **447.83** | **39.57** | **100.00** | **99.97** | **99.66** |
| **Marinobacter sp.LV10MA510-1** | **135.05** | **12.86** | **99.84** | **99.85** | **98.96** |
| Muricauda sp.ES.050 | 618.76 | 50.23 | 100.00 | 99.92 | 99.70 |
| Psychrobacter sp.LV10R520-6 | 425.47 | 39.30 | 99.86 | 99.34 | 98.34 |
| Cohaesibacter sp.ES.047 | 170.59 | 17.91 | 99.54 | 99.28 | 98.32 |
| Thioclava sp.ES.032 | 78.32 | 9.49 | 99.56 | 99.47 | 99.32 |
| Propionibacteriaceae bacterium | 31.90 | 4.31 | 100.00 | 99.99 | 99.97 |
| Micromonospora echinofusca | 18.19 | 3.69 | 99.66 | 99.53 | 99.23 |
| Micromonospora echinaurantiaca | 14.91 | 3.02 | 99.59 | 99.67 | 99.19 |

**Supplementary Table 4.** Coverage and average nucleotide identity (ANI) of strains in the NWc data set.

| Genomes | GenBank no | ANI (%) |
|---|---|---|
| Streptococcus_thermophilus_isolate_NWC_1_1 | CP029252.1 | 99.99 |
| Streptococcus_thermophilus_isolate_NWC_2_1 | CP031021.1 | |
| Lactobacillus_delbrueckii_isolate_NWC_1_2 | CP029250.1 | 99.24 |
| Lactobacillus_delbrueckii_isolate_NWC_2_2 | CP031023.1 | |
| Lactobacillus_helveticus_isolate_NWC_2_4 | CP031018.1 | 98.03 |
| Lactobacillus_helveticus_isolate_NWC_2_3 | CP031016.1 | |

**Supplementary Table 5.** NWCs ONT. The genome fraction of each individual strain in the NWCs data (Illumina and ONT). Present the impact of different sequencing coverage in distinct assembly methods.

| Assembly | Coverage (Illumina) | Coverage (ONT) | HyLight | OPERA-MS | Unicycler | hybridSPAdes |
|---|---|---|---|---|---|---|
| Streptococcus_thermophilus_isolate_NWC_1_1 | 56.29 | 84.08 | 99.94 | 98.03 | 50.11 | 83.49 |
| Streptococcus_thermophilus_isolate_NWC_2_1 | 55.07 | 75.20 | 99.99 | 92.40 | 89.88 | 76.19 |
| Lactobacillus_delbrueckii_isolate_NWC_1_2 | 39.38 | 25.68 | 96.33 | 90.10 | 92.46 | 75.85 |
| Lactobacillus_delbrueckii_isolate_NWC_2_2 | 35.13 | 38.77 | 96.59 | 89.41 | 80.89 | 24.91 |
| Lactobacillus_helveticus_isolate_NWC_2_4 | 17.59 | 221.79 | 98.09 | 93.00 | 82.22 | 63.47 |
| Lactobacillus_helveticus_isolate_NWC_2_3 | 10.27 | 90.80 | 90.85 | 77.00 | 55.63 | 32.22 |

**Supplementary Table 6.** NWCs PacBio. The genome fraction of each individual strain in the NWCs data (Illumina and PacBio). Present the impact of different sequencing coverage in distinct assembly methods.

| Assembly | Coverage (Illumina) | Coverage (PacBio) | HyLight | OPERA-MS | Unicycler | hybridSPAdes |
|---|---|---|---|---|---|---|
| Streptococcus_thermophilus_isolate_NWC_1_1 | 56.29 | 243.26 | 96.30 | 95.25 | 98.77 | 82.99 |
| Streptococcus_thermophilus_isolate_NWC_2_1 | 55.07 | 190.32 | 93.67 | 83.22 | 89.53 | 69.80 |
| Lactobacillus_delbrueckii_isolate_NWC_1_2 | 39.38 | 180.43 | 93.58 | 86.48 | 97.58 | 75.01 |
| Lactobacillus_delbrueckii_isolate_NWC_2_2 | 35.13 | 36.19 | 72.76 | 81.37 | 82.84 | 20.96 |
| Lactobacillus_helveticus_isolate_NWC_2_4 | 17.59 | 1.45 | 70.26 | 65.34 | 39.30 | 54.69 |
| Lactobacillus_helveticus_isolate_NWC_2_3 | 10.27 | 2.47 | 41.08 | 40.54 | 12.38 | 37.36 |

**Supplementary Table 7.** A comparison of assembly result quality is performed between HyLight and Strainberry. Indels/100 kbp: average number of insertion or deletion errors per 100,000 aligned bases. Mismatches/100 kbp = average number of mismatch errors per 100,000 aligned bases. Genome Fraction GF reflects how much of each of the strain-specific genomes is covered by contigs. N/100 kbp denotes the average number of uncalled bases (N's) per 100,000 bases in contigs. MC = fraction of misassembled contigs.

| Assembly | GF(%) | Indels/100kbp | Mismatches/100kbp | NGA50 | N/100 kbp | MC(%) |
|---|---|---|---|---|---|---|
| 3 salmonella | | | | | | |
| HyLight | 97.13 | 1.27 | 15.00 | 356582 | 0.00 | 0.22 |
| Strainberry | - | - | - | - | - | - |
| 20 strains | | | | | | |
| HyLight | 92.15 | 5.55 | 58.66 | 139730 | 0.00 | 0.33 |
| Strainberry | 78.43 | 8.48 | 26.26 | 95338 | 19.60 | 2.30 |
| 100 strains | | | | | | |
| HyLight | 93.86 | 11.68 | 55.68 | 163296 | 0.00 | 1.00 |
| Strainberry | 79.25 | 210.30 | 115.87 | 65706 | 68.05 | 4.84 |
| 210 strains | | | | | | |
| strain_hybrid | 90.16 | 17.69 | 52.78 | 128015 | 0.00 | 1.63 |
| Strainberry | 78.56 | 418.36 | 102.55 | 81842 | 41.72 | 4.66 |
| Bmock12 ONT | | | | | | |
| HyLight | 99.77 | 1.45 | 3.58 | 281944 | 0.00 | 3.59 |
| Strainberry | 67.60 | 705.34 | 264.57 | 688598 | 3.57 | 11.66 |
| Bmock12 PacBio | | | | | | |
| HyLight | 98.57 | 5.29 | 19.24 | 123823 | 0.00 | 7.62 |
| Strainberry | 62.50 | 272.41 | 33.48 | 72377 | 21.30 | 22.17 |
| NWC ONT | | | | | | |
| HyLight | 95.35 | 30.45 | 174.89 | 62800 | 0.00 | 9.37 |
| Strainberry | 91.69 | 764.14 | 193.84 | 141570 | 60.26 | 22.94 |
| NWC PacBio | | | | | | |
| HyLight | 78.94 | 84.42 | 219.74 | 22388 | 0.00 | 4.27 |
| Strainberry | 43.03 | 246.94 | 111.47 | - | 0.65 | 22.22 |

**Supplementary Table 8.** Running times of different assembly approaches.

| Methods | CPU time (h) | Peak Memory Usage (GB) |
|---|---|---|
| OPERA-MS | 2.09 | 1.23 |
| hybridSPAdes | 5.53 | 3.85 |
| HyLight | 7.01 | 15.99 |
| MetaPlatanus | 6.93 | 69.26 |
| Unicycler | 53.71 | 6.53 |
